# No evidence for three functionally specialized subregions in the subthalamic nucleus: A model-based 7 T fMRI study

**DOI:** 10.1101/2021.08.12.456040

**Authors:** Steven Miletić, Max C. Keuken, Martijn Mulder, Robert Trampel, Gilles de Hollander, Birte U. Forstmann

## Abstract

The subthalamic nucleus (STN) is a small, subcortical brain structure. It is a target for deep brain stimulation, an invasive treatment that reduces motor symptoms of Parkinson’s disease. Side effects of DBS are commonly explained using the tripartite model of STN organization, which proposes three functionally distinct subregions in the STN specialized in cognitive, limbic, and motor processing. However, evidence for the tripartite model exclusively comes from anatomical studies and functional studies using clinical patients. Here, we provide the first experimental tests of the tripartite model in healthy volunteers using ultra-high field 7 Tesla (T) functional magnetic resonance imaging (fMRI). 34 participants performed a random-dot motion decision-making task with a difficulty manipulation and a choice payoff manipulation aimed to differentially affect cognitive and limbic networks. Moreover, participants responded with their left and right index finger, differentially affecting motor networks. We analysed BOLD signal in three subregions of equal volume of the STN along the dorsolateral-ventromedial axis, identified using manually delineated high resolution anatomical images. Our results indicate that all segments responded equally to the experimental manipulations, and did not support the tripartite model.

## 1. Introduction

The subthalamic nucleus (STN) is a small, subcortical brain structure and a node of the basal ganglia (BG). It is a target for deep brain stimulation (DBS), which reduces the motor symptoms of Parkinson’s Disease (PD) by electrically stimulating the STN (Fasano and Lozano, 2015; Limousin et al., 1995; Lozano and Lipsman, 2013). However, DBS of the STN can lead to severe side effects such as cognitive decline, depression, and (hypo)mania (Christen et al., 2012; Groiss et al., 2009; Temel et al., 2006).

The so-called tripartite model of the STN (e.g., Haynes and Haber, 2013; Joel and Weiner, 1997; Parent and Hazrati, 1995) is used to explain these side effects, by proposing that the STN is subdivided in three parts with different functional roles (Temel et al., 2005a). These subdivisions are hypothesized to be connected to three cortical networks, which can be characterized as a ‘limbic’ network, a ‘cognitive’ network, and a ‘motor’ network. Consequently, the proposed subdivision of the STN also consists of limbic, cognitive, and motor parts, which can be found in the ventromedial, central, and dorsolateral parts of the STN. Assuming this tripartite division, misplacements of the electrode in the limbic or cognitive parts of the STN, rather than the targeted motor part, are thought to be the origin of non-motor side effects of DBS of the STN (Karachi et al., 2009; Temel et al., 2005a).

Although empirical studies provide indirect evidence that the tripartite model is plausible in the human brain (Greenhouse et al., 2013; Haynes and Haber, 2013; Lambert et al., 2012; Mallet et al., 2007), a systematic review of the neuroanatomical literature on the STN in the non-human primate shows little consistency in the number of subdivisions and their topological organization (Keuken et al., 2012). Moreover, to the best of our knowledge, there is no study that shows direct evidence in a healthy human population that different subparts of the STN are involved in different functions. Currently, the evidence for putative functional subdivisions in the STN comes nearly exclusively from neuroanatomical data (Haynes and Haber, 2013; Joel and Weiner, 1997; Parent and Hazrati, 1995), rather than from functional data acquired during the execution of tasks that require limbic, cognitive or motor processing (but see Greenhouse et al., 2013, 2011 for results of a cognitive paradigm in a clinical group).

Here, we test the tripartite hypotheses by imaging the STN during a perceptual decision-making task (Ball and Sekuler, 1982; Britten et al., 1993, 1992; Pilly and Seitz, 2009), which requires participants to choose whether a cloud of randomly moving dots moves, on average, to the left or right. To modulate the hypothesized cortico-basal ganglia networks, two manipulations were added to the task based on earlier literature (Mulder et al., 2014). The first manipulation aimed to evoke limbic processing by inducing response biases. On half of the trials, subjects were cued to one of two response options with a higher potential payoff. Response bias manipulations have been associated with a ‘limbic’ network including orbitofrontal cortex, hippocampus, and ventral striatum (Forstmann et al., 2010b; Keuken et al., 2014b; Mulder et al., 2012, 2014; Summerfield and Koechlin, 2010). The second manipulation altered cognitive processing, by changing the difficulty of the stimulus discrimination task, which has been shown to change the rate of evidence accumulation during decision making. On half of the trials, the coherence of the dots on the screen was relatively high (easy trials), whereas on the other trials the coherence was relatively low (hard trials). This manipulation has been shown to modulate activity in ‘cognitive’ frontal cortical areas such as the dorsolateral prefrontal cortex (Heekeren et al., 2004; Kaiser et al., 2007), as well as in the insula (Binder et al., 2004; Ho et al., 2009; Keuken et al., 2014b; Thielscher and Pessoa, 2007). Furthermore, in earlier work using electrophysiological recordings and UHF-fMRI, the STN has been shown to reflect a ‘conflict’ or ‘normalization’ signal (Bogacz and Gurney, 2007; Frank et al., 2015; Keuken et al., 2015) that should be inversely proportional to the similarity of two choice options. As a third experimental factor of interest, we tested for motor-related processing in the STN by analyzing response directions. The STN is considered part of a cortico-basal ganglia-thalamic ‘motor’ loop, including primary motor cortex (M1; Alexander and Crutcher, 1990; Temel et al., 2005a, 2005b). The lateralization of motor activity related to response hands in M1 forms a benchmark fMRI finding (Dassonville et al., 1997; Kim et al., 1993a, 1993b) and we hypothesized that such a lateralization might also occur in the STN (Devos et al., 2006).

While performing the task, participants were scanned using ultra-high field (UHF) 7 Tesla functional MRI. Unlike fMRI at lower fields, UHF fMRI can potentially resolve fine activation patterns within the STN, because of its increased spatial resolution and signal-to-noise ratios (de Hollander et al., 2017; Miletić et al., 2020; van der Zwaag et al., 2016). To maximize anatomical specificity of our functional measurements, we used manually delineated, anatomically defined masks of the STN, based on high resolution structural images (Keuken et al., 2014a). These STN masks were subdivided in three parts of equal volume along their dorsolateral-ventromedial axis (see Figure 1) using an automated procedure. To maximize the sensitivity of the functional imaging data, an optimized fMRI protocol was used that maximizes blood-oxygenation level dependent (BOLD) sensitivity in the STN (de Hollander et al., 2017; Miletić et al., 2020).

**Figure 1.**
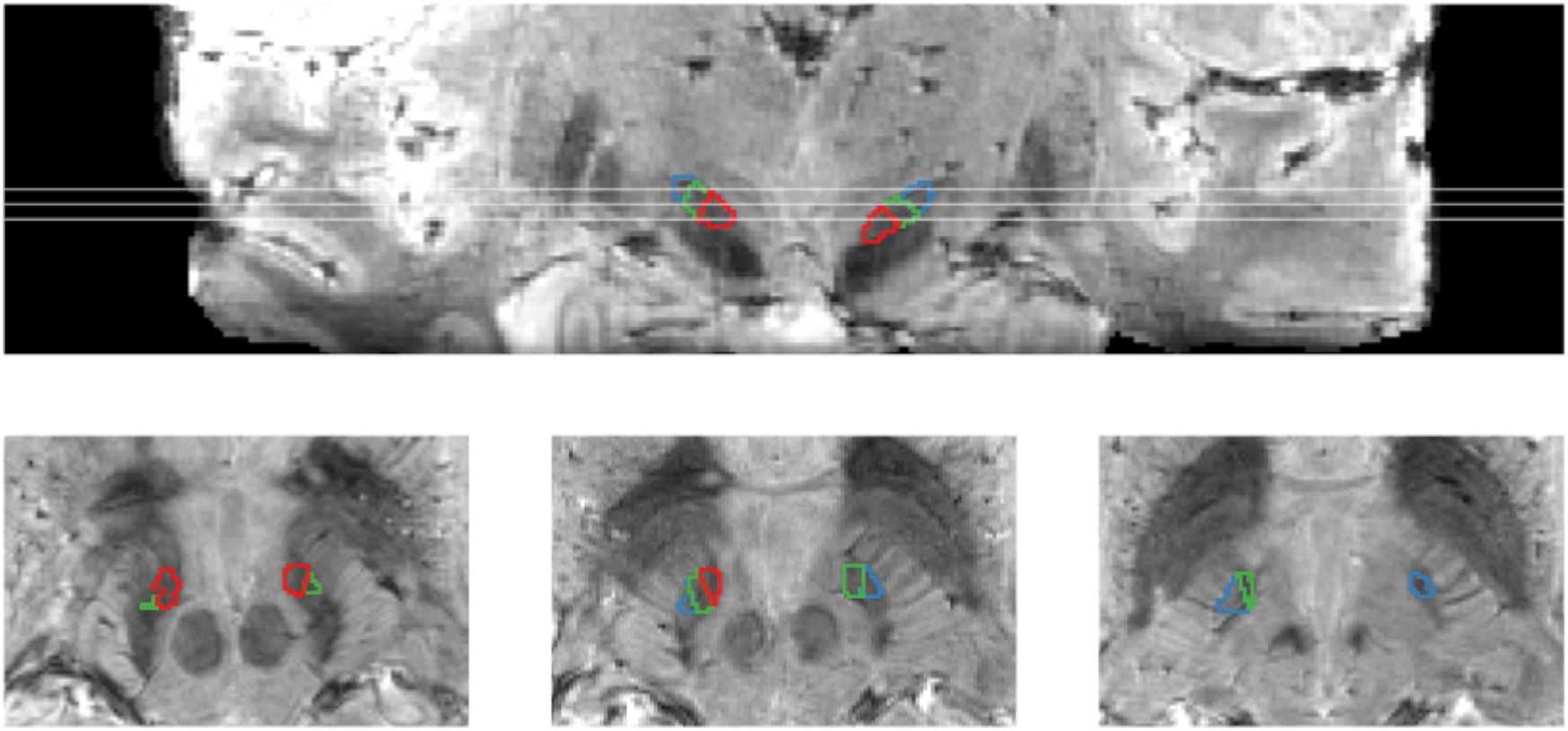
PCA segments in individual space overlaid on the 0.5 mm isotropic averaged ME-GRE image (neurological convention). Top row corresponds to a coronal view, where three white lines indicate the axial cuts illustrated on the bottom. The blue region corresponds to the dorsolateral segment A, the green region to central segment B, and the red region to ventromedial segment C.

If the STN has functional subdivisions that are relevant for perceptual decision making, these three segments should be differentially modulated by the three task factors. Specifically, the ventromedial ‘limbic’ segment would be most sensitive to the response bias cues, (b) the central ‘cognitive’ segment would be most sensitive to the difficulty manipulation, and (c) the dorsolateral ‘motor’ segment would be most sensitive to the response hand.

To test these hypotheses, we first contrasted the BOLD responses during the main task conditions. Specifically, we tested for differences between BOLD responses between payoff and neutral cues (limbic), between easy and hard stimuli (cognitive), and between the two response hands (motor). Furthermore, we tested whether these contrasts had different sizes in the three STN subdivisions. Subsequently, we adopted a model-based cognitive neuroscience approach, capitalizing on individual differences. This approach potentially increases sensitivity and specificity to differential activation patterns (de Hollander et al., 2016; Forstmann et al., 2017, 2011; Turner et al., 2019, 2017). Concretely, we fit the diffusion decision model (DDM; Forstmann et al., 2016; Ratcliff, 2006, 1978; Ratcliff et al., 2016; Ratcliff and McKoon, 2008) to the behavioral data. We then tested whether individual differences in behavioral performance, as quantified by the DDM, covaried with the individual differences in the size of the corresponding fMRI contrasts (Forstmann et al., 2010b, 2008; Mulder et al., 2014, 2012). In this way, we tested the hypothesis that the different segments were specifically related to the two latent processes underlying the two task manipulations.

## 2. Methods

### 2.1 Participants

We analyzed two data sets collected at different scanner sites: Max Planck Institute for Human Brain and Cognitive Sciences in Leipzig, Germany, and the Spinoza Centre for Neuroimaging in Amsterdam, the Netherlands. The behavioral paradigms were identical but for one task parameter, and data acquisition protocols were built to be as identical as possible within the technical constraints of both scanners.

In dataset 1, 19 healthy subjects were scanned (10 males; mean age 26.9, std. age 2.4, range 23 – 32). All subjects had normal or corrected to normal vision and no history of neurological or psychological disorders. All subjects were right-handed, as confirmed by the Edinburgh Inventory (Oldfield, 1971). All subjects participated in an earlier study using both structural and functional MRI in the basal ganglia (de Hollander et al., 2017). The study was approved by the ethical committee at the Medical Faculty, Leipzig University, Germany. All subjects gave written informed consent and received a monetary reward for their participation, as well as an additional monetary reward based on their task performance.

In dataset 2, 14 healthy subjects were scanned (7 males; mean age 24.14, std. age 3.3, range 20 - 29). Two participants were left-handed. The study was approved by the local ethical committee of the University of Amsterdam, the Netherlands. All subjects gave written informed consent and received a monetary reward for their participation, as well as an additional monetary reward based on their task performance.

### 2.2 Experimental paradigm

The experimental paradigm is illustrated in Figure 2A. The task paradigm was a random-dot motion (RDM) task (Ball and Sekuler, 1982; Britten et al., 1993, 1992; Pilly and Seitz, 2009), in which the participant is presented with a cloud of randomly moving dots. A subset of the dots, determined by the coherence level, moves coherently to the left or right. The participant is instructed to decide in which direction the cloud moves on average by making a button press with the left or right hand. Response time is defined as the time between the onset of the RDM stimulus and the button press.

**Figure 2.**
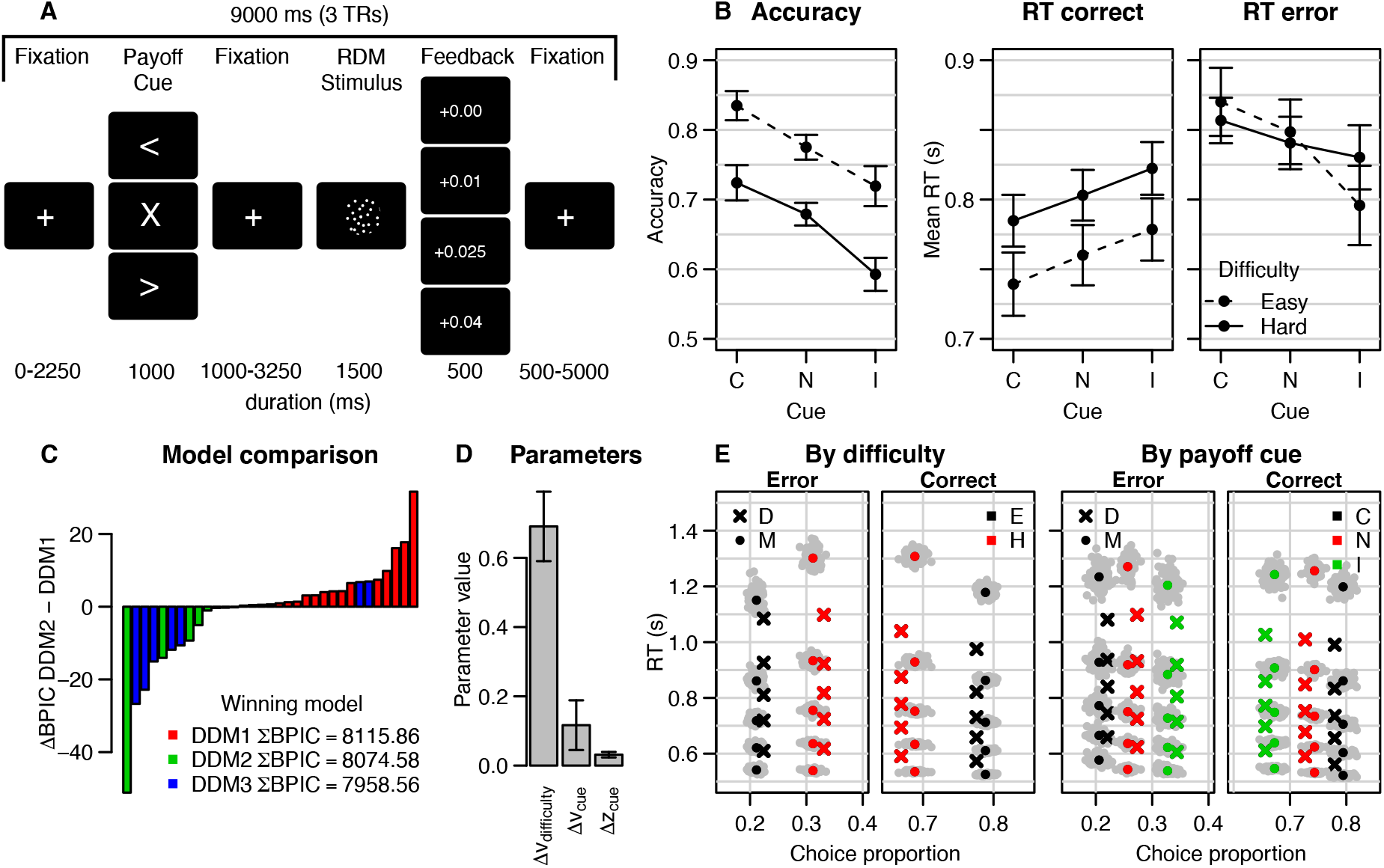
A. Example trial of the experimental paradigm. Each trial started with a fixation cross, followed by the potential payoff cue, which was either an arrow pointing leftwards or rightwards (biased trials) or an X (neutral trials). After the potential payoff cue, the fixation cross was shown again, followed by the random-dot motion stimulus, during which the participants made their response by means of a button press. After 1.5 s, feedback was shown to present the reward earned, which depended on the accuracy of the response and the congruency of the potential payoff cue. The trial ended with a fixation cross. B. Behavioral data. Error bars are within-subject standard error (Cousineau, 2005). Labels on the x-axis are congruent (C), neutral (N), or incongruent (I) potential payoff cue conditions. C. Model comparison. The bar plot shows, per subject, the difference between BPIC-values for DDM1 (starting point shift) and DDM2 (drift rate shift), to compare the strength of evidence for one over the other strategy. Colors indicate winning model per participant; the overall winning model was DDM3 (lowest summed BPIC) with *both* drift rate and starting point varying by potential payoff cue. D. Size of parameter changes due to the experimental manipulations. v = drift rate, z = starting point. E. Quantile probability plot of model fit (model 2). Crosses are data (D), circles are model (M), colors indicate the difficulty (left two panels, easy (E) and hard (H)) or congruency of the potential payoff cue (congruent (C), neutral (N), incongruent (I)). Grey dots are individual predictions using a random sample from the posteriors, the colored circles are the mean of these posteriors predictive data points. The model overestimates the skewness of the RT distributions, which combined with the slow errors in the data, is suggestive of an urgency signal. The model furthermore overestimates accuracy after incongruent potential payoff cues.

We used a factorial design with potential payoff cue type and RDM stimulus difficulty as independent variables. Every trial (384 in total) started with a potential payoff cue (one out of three different options), followed by the RDM stimulus (one out of two different difficulty levels), and feedback. The potential payoff cues consisted of an arrow pointing to the left (25% of the trials) or right (25% of the trials), or a cross (’neutral’ condition, the remaining 50% of the trials). Subjects could earn additional monetary reward based on their performance. The potential payoff cue indicated different *potential monetary rewards:* different responses yielded different payoffs, provided that the response was correct. Specifically, if a response was correct *and* congruent with the direction of the potential payoff cue arrow, the subject earned 0.04 euro for that trial. However, if the response was correct, but incongruent with the potential payoff cue arrow, the subject earned only 0.01 euro for that trial. If a response was correct on a neutral trial, the subject earned 0.025 euro. When subjects gave the incorrect response, they did not gain any reward, regardless of the cued direction.

The RDM consisted of a cloud of dots in a circle with a diameter of 5 degrees (dva) and with on average 16.7 dots per square degree. The dots moved around with a speed of 5 dva per second. Frames were presented at a speed of 60 frames per second. On the first three frames, the dots were randomly positioned within the circle. The task then subsequently looped over three frames while the dots of the presented frame were repositioned. Before a frame was drawn, a portion of dots, determined by the ‘coherence’ parameter (determining the difficulty of the trial), was repositioned a fixed amount to the left or right (making sure that the speed was 5 degrees per second). The remaining portion of dots was moved by the same amount, but in a random direction. The RDM was always presented for 1500 ms, independent of the subject responses, to prevent visual stimulus duration from confounding response behavior. In dataset 1, the coherences were set to 16% (easy, 50% of trials) and 8% (hard, 50% of trials). In dataset 2, the coherences were set to 35% (easy, 50% of trials) and 15% (hard, 50% of trials). Since the analyses below rely on within-subject *differences* in behavior between difficulty conditions (and no ceiling or floor performance was reached), this between-dataset difference in the coherence levels should not affect the results. Regardless, we accounted for any potential confounds by incorporating a random effect of dataset in the whole-brain GLM to model any dataset-related effects in the group statistics. This random effect was not significant in any of the regions of interest for any of the contrasts of interest.

After the RDM stimulus, feedback was presented on whether they were correct and how much money they earned on that trial ‘+€0.01’ (incongruent potential payoff cue), ‘+€0.025’ (neutral potential payoff cue), or ‘+€0.04’ (congruent potential payoff cue) for correct trials, or ‘+€0.00’ for incorrect trials. If subjects responded in less than 250 ms, or needed more than 1250 ms, they received the feedback ‘too fast’ or ‘too slow’, respectively.

### 2.3 General Procedure

After the participants were screened for MRI, they were introduced to the task on a laptop. They were explicitly explained how the cue payoffs determine the amount of money earned per trial, with a few examples. The participants were then also explained why, with limited information, following the potential payoff cues is a rational strategy. After this introduction, subjects performed 384 trials of an RDM task with potential payoff cues and two difficulty levels, divided over 3 runs, each consisting of 128 trials in 19 minutes and 21 seconds (approximately one hour of functional scanning in total). Over these 384 trials, subjects could gain an additional monetary reward, based on their performance, of at most 9 euro and 60 cents. A trial always took 9 seconds (3 TRs) in total (see Figure 2A). It started with a fixation cross for 0, 750, 1500, or 2250 ms (pseudo-randomly sampled), after which the response payoff cue was presented for 1000 ms. After the potential payoff cue, a second fixation cross was presented for 1000, 1750, 2500, or 3250 ms. Subsequently, the dots were presented for 1500 ms and the subject had to respond with their left or right index finger. Immediately after the RDM stimulus, feedback was presented for 500 ms. It showed how much the subject earned on that trial. For the remaining 500 – 5000 ms in each trial, a blank screen was presented.

### 2.4 MRI scanning protocols

Functional neuroimaging of the basal ganglia at UHF is challenging compared to most cortical brain regions (de Hollander et al., 2015; Forstmann et al., 2017; Keuken et al., 2018a). Therefore, we drew upon earlier validated MRI protocols for both structural (Keuken et al., 2014a) and functional (de Hollander et al., 2017; Miletić et al., 2020) imaging of the STN. Specifically, in dataset 1, subjects were scanned in a Siemens MAGNETOM 7 T system (Siemens Healthineers, Germany) with a 32 channel phased array head coil (Nova Medical Inc, USA). We used T2*-weighted images from a spoiled multi-echo Gradient-Recalled Echo (ME-GRE) protocol (TEs = 11.22, 20.39, 29.57 ms) with a resolution of 0.5 mm isotropic that were collected during earlier studies (de Hollander et al., 2017; Keuken et al., 2014a) for visualization of the STN and its neighboring structures. We also acquired 0.7 mm isotropic T1-weighted images using a MP2RAGE sequence (Marques et al., 2010) for registration purposes.

For the functional imaging, we used a recently developed BOLD fMRI protocol that balances spatial resolution with subcortical SNR and has a relatively short echo time, accounting for the relatively very short T_2_*-relaxation values in the STN (Aquino et al., 2009; de Hollander et al., 2017; Deistung et al., 2013; Keuken et al., 2017, 2014a). In dataset 1, a T_2_*-weighted 2D-EPI protocol with a spatial resolution of 1.5 mm isotropic, TR = 3000 ms, slice thickness 1.5 mm, 90 slices, interleaved acquisition, TE = 14 ms, flip angle = 60°, bandwidth 1446 Hz/Px, echo spacing 0.8 ms, FOV 192 × 192 × 135 mm, phase encoding direction A ≫ P, partial Fourier 6/8, GRAPPA acceleration factor 3, matrix size 128 × 128 was used. Due to the increased number of slices compared to the protocol presented in de Hollander et al. (2017; 90 vs. 60), the acquired volume now covered the entire brain for all subjects. To correct for field inhomogeneities, a corresponding B0 field map with the same FOV was acquired (TR = 1500 ms, TE_1-2_ = 6, 7.02 ms).

In dataset 2, participants were scanned in a Philips Achieva 7 T system (Philips, The Netherlands). The anatomical images of these subjects were already collected as part of a database (Alkemade et al., 2020), in which an MP2RAGE-ME protocol (Caan et al., 2019) was used (0.7 mm isotropic) that was used to calculate quantified T1, T_2_*, and QSM images. For functional imaging, we used a T_2_*-weighted 2D-EPI protocol with a resolution of 1.5 mm isotropic, TR = 3000 ms, slice thickness 1.5 mm, 80 slices, interleaved acquisition, TE = 14 ms, flip angle = 60°, bandwidth 1461 Hz/Px, echo spacing 0.85 ms, FOV 192 × 192 × 120 mm, phase encoding direction A ≫ P, HalfScan 0.76, SENSE acceleration factor 3, matrix size 128 × 128 (since the gradients have a lower slew rate, to keep TRs consistent, we could acquire less slices than in dataset 1). Again, we collected a B0 field map with the same FOV.

We would like to note that due to the heavy-duty gradient cycles in our fMRI protocols, there is the potential for serious heating of the gradient coils. Indeed, when running pilots for follow-up studies using the same sequence at the Philips site (dataset 2), we noticed that the temperature in the bore notably increased during scanning. The scanner system never reported temperatures outside its safety limits, but we would like to urge researchers to exhaustively pilot new studies with gradient-heavy fMRI protocols such as ours on a phantom, as well as monitor the development of the temperature inside the bore and on the bore wall during a multi-run scan protocol, before commencing data collection on human subjects, and always ask about the temperature in the bore during subject debriefings.

### 2.5 Anatomical Labeling

In dataset 1, the left and right STN were manually delineated on the 0.5 mm isotropic T_2_*-weighted ME-GRE images by two independent raters, following a previously published protocol (de Hollander et al., 2017; Keuken et al., 2014a). Similar to earlier studies, only voxels that were labeled as part of the STN by both raters were included in further analyses. In dataset 2, the manual delineations were based on QSM images obtained from the MP2RAGE-ME (Caan et al., 2019) sequence. The database included manual delineations by two independent rates. Again, only voxels that were labeled as part in the STN by both raters were included in the analyses.

To increase the limited functional signal-to-noise ratio in the STN without resorting to spatial smoothing, we subdivided the STN in smaller segments a-priori. All individual STN masks were transformed to the space of the ICBM 152 Nonlinear Asymmetrical template version 2009c (MNI2009c; Fonov et al., 2009). On the mean-subtracted coordinates of the voxels in these masks, we performed a principal components analysis to find the 3D vector (direction) that explained the most variance in the coordinates (Alkemade et al., 2019). We interpreted this vector as the ‘main ventromedial-dorsolateral’ axis of the STN (de Hollander et al., 2014; Haynes and Haber, 2013; Temel et al., 2005a).

Then, for every individual subject mask *separately*, all voxel coordinates on this axis were given a ‘ventromedial score’. This voxel-wise score allowed us to divide individual left and right STN masks into three segments: the ‘posterior-dorsolateral’ (segment A), the ‘central’ segment (segment B), and the ‘anterior-ventromedial’ segment (segment C; see Fig. 1). Note that these segments always covered the entire individual anatomical mask for each subject separately, with an equal volume. The segments were transformed to individual functional (EPI) space by the inverse transformation from individual to standard MNI space, as provided by the fmriprep registration procedure, using a single interpolation step, and used as regions of interest (ROIs) in further analyses. In the individual functional space at 1.5 mm isotropic resolution, the segments had a volume of on average 6.1 voxels (20.63 mm^3^; std. 2.9 voxels, 9.66 mm^3^) and never overlapped with other segments.

### 2.6 MRI data preprocessing

Results included in this manuscript come from preprocessing performed using *fMRIPrep* 20.0.0 (Esteban et al., 2019, 2018; RRID:SCR_016216), which is based on *Nipype* 1.4.2 (Gorgolewski et al., 2018 RRID:SCR_002502, 2011).

#### 2.6.1 Anatomical data preprocessing

The T1-weighted (T1w) image was corrected for intensity non-uniformity (INU) with N4BiasFieldCorrection (Tustison et al., 2010), distributed with ANTs 2.2.0 (Avants et al., 2008 RRID:SCR_004757), and used as T1w-reference throughout the workflow. The T1w-reference was then skull-stripped with a *Nipype* implementation of the antsBrainExtraction.sh workflow (from ANTs), using OASIS30ANTs as target template. Brain tissue segmentation of cerebrospinal fluid (CSF), white-matter (WM) and gray-matter (GM) was performed on the brain-extracted T1w using fast (FSL 5.0.9, RRID:SCR_002823; Zhang et al., 2001). Brain surfaces were reconstructed using recon-all (FreeSurfer 6.0.1, RRID:SCR_001847; Dale et al., 1999), and the brain mask estimated previously was refined with a custom variation of the method to reconcile ANTs-derived and FreeSurfer-derived segmentations of the cortical gray-matter of Mindboggle (RRID:SCR_002438; Klein et al., 2017). Volume-based spatial normalization to one standard space (MNI152NLin2009cAsym) was performed through nonlinear registration with antsRegistration (ANTs 2.2.0), using brain-extracted versions of both T1w reference and the T1w template. The following template was selected for spatial normalization: *ICBM 152 Nonlinear Asymmetrical template version 2009c* (Fonov et al., 2009; RRID:SCR_008796; TemplateFlow ID: MNI152NLin2009cAsym).

#### 2.6.2 Functional data preprocessing

For each of the 3 BOLD runs found per subject (across all tasks and sessions), the following preprocessing was performed. First, a reference volume and its skull-stripped version were generated using a custom methodology of *fMRIPrep*. In dataset 1, a B0-nonuniformity map (or *fieldmap*) was estimated based on a phase-difference map calculated with a dual-echo GRE (gradient-recall echo) sequence, processed with a custom workflow of SDCFlows inspired by the epidewarp.fsl script and further improvements in HCP Pipelines (Glasser et al., 2013). In dataset 2, a B0-nonuniformity map was directly measured with an MRI scheme designed with that purpose (typically, a spiral pulse sequence). The *fieldmap* was then co-registered to the target EPI (echo-planar imaging) reference run and converted to a displacements field map (amenable to registration tools such as ANTs) with FSL’s fugue and other *SDCflows* tools. For one subject of dataset 2, no B0-nonuniformity map was available. For this subject, a deformation field to correct for susceptibility distortions was estimated based on fMRIPrep’s fieldmap-less approach. The deformation field is that resulting from co-registering the BOLD reference to the same-subject T1w-reference with its intensity inverted (Huntenburg, 2014; Wang et al., 2017). Registration is performed with antsRegistration (ANTs 2.2.0), and the process regularized by constraining deformation to be nonzero only along the phase-encoding direction, and modulated with an average fieldmap template (Treiber et al., 2016). Based on the estimated susceptibility distortion, a corrected EPI (echo-planar imaging) reference was calculated for a more accurate co-registration with the anatomical reference. The BOLD reference was then co-registered to the T1w reference using bbregister (FreeSurfer) which implements boundary-based registration (Greve and Fischl, 2009). Co-registration was configured with six degrees of freedom. Head-motion parameters with respect to the BOLD reference (transformation matrices, and six corresponding rotation and translation parameters) are estimated before any spatiotemporal filtering using mcflirt (FSL 5.0.9; Jenkinson et al., 2002). BOLD runs were slice-time corrected using 3dTshift from AFNI 20160207 (Cox and Hyde, 1997; RRID:SCR_005927). The BOLD time-series (including slice-timing correction when applied) were resampled onto their original, native space by applying a single, composite transform to correct for head-motion and susceptibility distortions. These resampled BOLD time-series will be referred to as *preprocessed BOLD in original space*, or just *preprocessed BOLD*. The BOLD time-series were resampled into standard space, generating a *preprocessed BOLD run in MNI152NLin2009cAsym space*. First, a reference volume and its skull-stripped version were generated using a custom methodology of *fMRIPrep*. Several confounding time-series were calculated based on the *preprocessed BOLD*: framewise displacement (FD), DVARS and three region-wise global signals. FD and DVARS are calculated for each functional run, both using their implementations in *Nipype* (following the definitions by Power et al., 2014). The three global signals are extracted within the CSF, the WM, and the whole-brain masks. Additionally, a set of physiological regressors were extracted to allow for component-based noise correction (CompCor; Behzadi et al., 2007). Principal components are estimated after high-pass filtering the *preprocessed BOLD* time-series (using a discrete cosine filter with 128s cut-off) for the two *CompCor* variants: temporal (tCompCor) and anatomical (aCompCor). tCompCor components are then calculated from the top 5% variable voxels within a mask covering the subcortical regions. This subcortical mask is obtained by heavily eroding the brain mask, which ensures it does not include cortical GM regions. For aCompCor, components are calculated within the intersection of the aforementioned mask and the union of CSF and WM masks calculated in T1w space, after their projection to the native space of each functional run (using the inverse BOLD-to-T1w transformation). Components are also calculated separately within the WM and CSF masks. For each CompCor decomposition, the *k* components with the largest singular values are retained, such that the retained components’ time series are sufficient to explain 50 percent of variance across the nuisance mask (CSF, WM, combined, or temporal). The remaining components are dropped from consideration. The head-motion estimates calculated in the correction step were also placed within the corresponding confounds file. The confound time series derived from head motion estimates and global signals were expanded with the inclusion of temporal derivatives and quadratic terms for each (Satterthwaite et al., 2013). Frames that exceeded a threshold of 0.5 mm FD or 1.5 standardised DVARS were annotated as motion outliers. All resamplings can be performed with *a single interpolation step* by composing all the pertinent transformations (i.e. head-motion transform matrices, susceptibility distortion correction when available, and co-registrations to anatomical and output spaces). Gridded (volumetric) resamplings were performed using antsApplyTransforms (ANTs), configured with Lanczos interpolation to minimize the smoothing effects of other kernels (Lanczos, 1964). Non-gridded (surface) resamplings were performed using mri_vol2surf (FreeSurfer).

Many internal operations of *fMRIPrep* use *Nilearn* 0.6.1 (Abraham et al., 2014; RRID:SCR_001362), mostly within the functional processing workflow. For more details of the pipeline, see the section corresponding to workflows in *fMRIPrep*’s documentation (https://fmriprep.readthedocs.io/en/latest/workflows.html).

### 2.7 Data analysis

Two subjects performed only 256, instead of 384 trials, due to fatigue (the entire experiment took approximately an hour). After careful inspection of the data set, these subjects were included in the analyses to improve statistical power.

#### 2.7.1 Behavioral analyses

To test whether the experimental manipulations were successful, behavioral data were analyzed using frequentist mixed effects models (MEMs; Barr et al., 2013; Gelman and Hill, 2007). Two MEMs were fit: One logistic model, to test the effects of the two manipulations on choice accuracy, and one linear model, to test the effects on choice response times (RTs). The logistic MEM included two main task manipulations: potential payoff cue congruency (congruent, neutral, incongruent), and difficulty (easy, hard), as well as their interaction. The linear MEM included the aforementioned task manipulations as well as the response accuracy, plus all interactions. For both models, subjects were included as random intercepts. We report estimates of the MEMs coefficients, their standard errors (SE), 95% confidence intervals (CIs), and a *p*-value obtained from a *t*-test using Satterthwaite’s method to estimate the degrees of freedom (Satterthwaite, 1941). These analyses made use of software packages *lme4* (Bates et al., 2015) and *lmerTest* (Kuznetsova et al., 2017) for the *R* programming language (R Core Team, 2017).

Next, we used the DDM to estimate parameters quantifying key properties of the latent cognitive processes underlying decision making. The DDM assumes that participants gradually accumulate noisy evidence for each choice option, until a threshold level of evidence is reached. At that point, the participant commits to the decision and executes the motor response. The process is characterized by four main parameters: The starting point of evidence accumulation *z*, which lies between thresholds *-a* and *a* which quantifies response caution, the mean speed of evidence accumulation *v* (the drift rate), and the non-decision time *t0* that quantifies the time required for perceptual encoding processes and motor execution. The ‘full’ DDM assumes additional parameters for between-trial variabilities in drift rate *s*_*v*_, starting point *s*_*z*_ and non-decision time *s*_*t*0_, which we did not estimate but fixed to 0. The between-trial variabilities in starting point and drift rate have been shown to be difficult to accurately recover (Boehm et al., 2018a). However, in an exploratory analysis, we also fit the winning DDM (reported below) including *s*_*v*_ and *s*_*z*_, and found that the estimates of *v*, *a*, *z*, and *t*0 were highly correlated (all *r* > 0.99) with the same estimated obtained from a DDM with *s*_*v*_ and *s*_*z*_ constrained to 0. Hence, not estimating *s*_*v*_ and *s*_*z*_ did not bias our parameters (for further discussion on estimating the between-trial variability parameters, see Boehm et al., 2018a). Within-trial noise parameter *s* was fixed to 1 to satisfy scaling constraints (Donkin et al., 2009; van Maanen and Miletić, 2020).

We fit three different DDMs to the behavioral data. All three DDMs assumed that choice difficulty affected drift rates (e.g., Palmer et al., 2005; Ratcliff and McKoon, 2008; Voss et al., 2004), which we modelled by independently estimating two drift rates, one per difficulty condition. The DDMs differed in how they explained the effect of the payoff cues: DDM1 assumed potential payoff cues biased starting points *z* (Diederich and Busemeyer, 2006; Edwards, 1965; Link and Heath, 1975; Mulder et al., 2012; Ratcliff, 1985; Ratcliff and Rouder, 1998; Voss et al., 2004); DDM2 assumed potential payoff cues biased drift rates (Ashby, 1983; Diederich and Busemeyer, 2006; Mulder et al., 2012; Ratcliff, 1981; Urai et al., 2019); and DDM3 assumed both types of bias were present (Mulder et al., 2012).

To model starting point bias, we estimated the neutral starting point plus a starting point shift in the direction of the potential payoff cue, such that *z*_*left cue*_ = *z*_*neutral*_ + *z*_*cue−shift*_ and *z*_*right cue*_ = *z*_*neutral*_ − *z*_*cue−shift*_ (in our parametrization, the DDM’s upper threshold corresponds to the leftward response). Similarly, to model drift rate bias, we assumed that *v*_*left cue*_ = *v* + *v*_*cue−shift*_ and *v*_*right cue*_ = *v* − *v*_*cue−shift*_, where the sign of *v* is negative if the dots move towards the right (and the size of *v* depends on the difficulty condition). In total, the three DDMs had six (*v*_*easy*_, *v*_*hard*_, *z*_*neutral*_, *z*_*cue−shift*_, *a*, *t*0), six (*v*_*easy*_, *v*_*hard*_, *v*_*cue−shift*_, *z*, *a*, *t*0), and seven (*v*_*easy*_, *v*_*hard*_, *v*_*cue−shift*_, *z*_*neutral*_, *z*_*cue−shift*_, *a*, *t*0) free parameters, respectively.

Since we intend to use the median posterior parameter estimates in correlation analyses, we did not estimate parameters hierarchically, since this would create a dependency between subject-level parameters and violate the assumption of independence of observations in correlation tests (Boehm et al., 2018b). Subject-level posterior distributions of each model’s parameters were estimated using a combination of differential evolution (DE) and Markov-chain Monte Carlo sampling (MCMC) with Metropolis-Hastings (Ter Braak, 2006; Turner et al., 2013). The default sampling settings, as implemented in the Dynamic Models of Choice R software (Heathcote et al., 2019), were used: The number of chains, D, was three times the number of free parameters. Cross-over probability (as part of the differential evolution) was set to 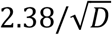. Migration probability was set to 0.05 (only during burn-in). Convergence was assessed using visual inspection of the chain traces and Gelman-Rubin diagnostic (Brooks and Gelman, 1998; Gelman and Rubin, 1992) (individual and multivariate potential scale factors < 1.02 in all cases). Datapoints with responses faster than 150 ms were considered as guesses and removed prior to fitting (cf. Ratcliff and Childers, 2015; Ratcliff and Tuerlinckx, 2002).

Independent truncated normal distributions were used as mildly informed priors for each parameter, as follows: *a*~*N*(2,0.5) truncated at 0 and 5, *v*_*easy/hard*_~*N*(2,0.5) truncated at −5 and 7, *z*~*N*(0.5,0.1), truncated at 0.3 and 0.7, *t*0~*N*(0.3,0.05) truncated at 0 and 1 s, *v*_*cue−shift*_~*N*(0,1) truncated at −5 and 7, and *z*_*cue−shift*_~*N*(0,0.1) truncated at −0.3 and 0.3.

The three DDMs were compared using the Bayesian Predictive Information Criterion (BPIC; Ando, 2007). Lower BPIC values indicate a better trade-off between quality of fit and model complexity. To visualize the quality of model fit of the winning model, we took 100 random samples from the posterior parameter distributions, with which we simulated the DDM (using the same trial numbers as in the experiment). We then calculated the 10^th^, 30^th^, 50^th^, 70^th^ and 90^th^ simulated RT quantile per condition for both the correct and error responses, and compared their (across-simulation) mean values with the empirical data (Figure 2E).

#### 2.7.2 Whole-brain GLMs

We then fit voxel-wise whole-brain GLMs to the fMRI data, in order to test whether the whole-brain BOLD contrasts are in line with the literature. In the first-level GLMs, we included eight task regressors: Leftward, rightward, and neutral potential payoff cues (on cue onset); easy and hard stimuli (on RDM stimulus onset); left and right responses (on response time); and errors (on feedback onset). Errors were explicitly modeled to decorrelate the effects of task difficulty from the effects of error (feedback) processing. It is well-known that limbic structures show heightened activity after an error has been made (e.g., Alexander and Brown, 2011; Klein et al., 2013; Ullsperger et al., 2010). Therefore, ‘drift rate’-related activity in limbic areas such as the insula might be a result of a larger number of errors in trials with a lower drift rate, unless the BOLD response related to errors is modeled. All eight task regressors were convolved in the canonical double-gamma hemodynamic response function (HRF; Glover, 1999), and their temporal derivatives were included in the design matrix. Additionally, we included a set of 16 cosines to model low-frequency drifts, and seven head-motion parameters (translation and rotation in 3 dimensions, and framewise displacement), DVARS (Power et al., 2014), and the first five aCompCor components (Behzadi et al., 2007) to model physiological noise. GLMs were fit using FSL FEAT (Woolrich et al., 2001). A smoothing kernel (FWHM = 5 mm, c.f. de Hollander et al., 2017; Miletić et al., 2020) was used to spatially smooth the data prior to fitting (Smith and Brady, 1997).

In a second-level GLM, the three runs per subject were combined using a fixed effects analysis. In a third-level GLM, FSL’s FLAME1 (Woolrich et al., 2004) was used. The design matrix included a group-wise intercept, a dummy variable to model potential dataset differences, and three model-based subject-wise covariates: (1) the drift rate difference (*v*_*hard*_ − *v*_*easy*_); (2) the starting point shift (*z*_*cue−shift*_); (3) the drift rate shift (*v*_*cue−shift*_). These covariates were demeaned.

Three main contrasts of interest were defined: (1) Response left – response right, to test for motor lateralization effects; (2) Payoff – neutral cues, to test for group-level main effects (i.e., testing for regions showing differential BOLD-responses after payoff compared to neutral cues) as well as covariances with potential payoff cue-related starting point or drift rate shifts (i.e., testing for regions in which the BOLD contrast size covaries with *z*_*cue−shift*_ or *v*_*cue−shift*_); (3) Hard – easy stimuli, to test for group-level main effects and across-subject covariances with drift rate shifts (*v*_*hard*_ − *v*_*easy*_). All third-level statistical parametric maps (SPMs; Figure 3) were thresholded using FSL’s *cluster* by first voxel-wise thresholding at *z* = 3.1 and then at *p* < 0.05 at the cluster level (Worsley, 2001).

**Figure 3.**
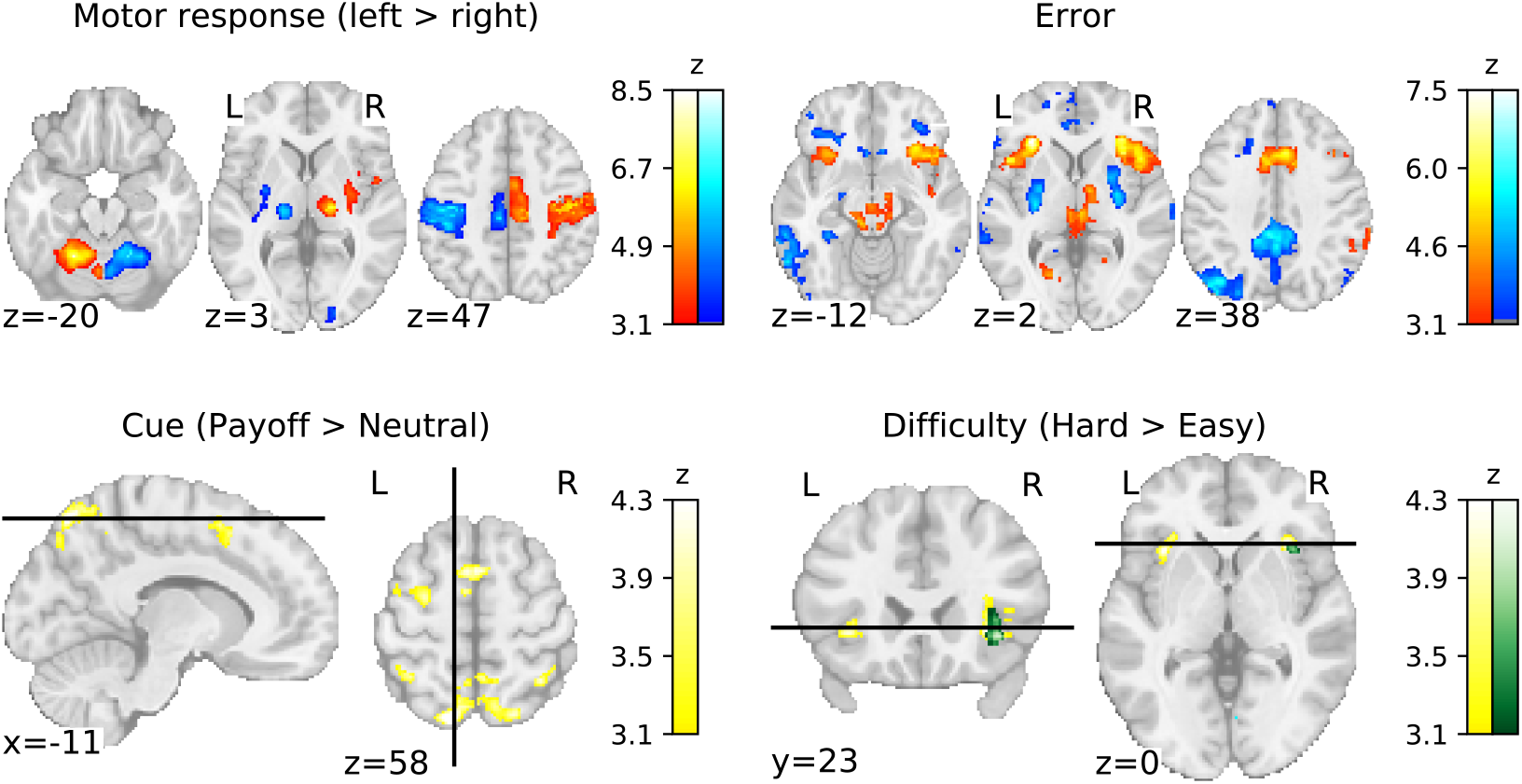
Statistical parametric maps of the main contrasts of interest in the whole-brain voxel-wise GLM analyses. Colors represent *z*-values of the group-level GLM, thresholded at *z* = 3.1 and *p* < 0.05 (two-sided), corrected at the cluster level using FSL’s family-wise error rate correction. Dual color bars are on the same scale, blue is used (top row) indicate negative *z*-values. For the difficulty contrast (bottom-left), yellow indicates the *z*-value of the group-level intercept, whereas the green indicates the *z*-values of the group-level covariance with the drift rate shift. Coordinates are in MNI 2009c (1mm) space.

#### 2.7.3 BOLD response deconvolutions

Next, we tested whether we had sufficient BOLD sensitivity, by estimating the shape and size of BOLD responses in the STN segments using deconvolution analyses (Dale, 1999; Gitelman et al., 2003; Glover, 1999; Poldrack et al., 2011). We first extracted the (unsmoothed) mean timeseries within each of the six (three per hemisphere) STN segments. Each timeseries was rescaled to percent signal change by dividing the timeseries by the mean signal value, multiplying by 100, and subtracting 100. The timeseries from the three runs per subject were then concatenated. Then, we performed a deconvolution using Fourier basis sets (Bullmore et al., 1996; Gitelman et al., 2003) with 7 regressors (an intercept, three sines, and three cosines) for two events of interest: The moving dots stimulus and potential payoff cue. Since the purpose of this deconvolution analysis is not to identify contrasts between the task manipulations, but instead to characterise the shape of the HRF, we ignored RDM stimulus difficulty and potential payoff cue types. Additionally, the model contained the same set of nuisance variables as used in the whole-brain GLM above: Five aCompCor components, 16 cosines to model low-frequency scanner drift, the DVARS, and seven head motion related parameters (rotation and translation in three dimensions, and the framewise displacement). The model was fit using ordinary least squares. Deconvolution analyses were performed using *nideconv* (de Hollander et al., 2019).

#### 2.7.4 ROI-wise GLMs

Subsequently, we defined four GLMs, each fit to the same six STN mean timeseries separately, to test how strongly the six segments responded to the task events. Importantly, and contrary to the whole-brain GLMs above, all STN timeseries were extracted from the unsmoothed functional timeseries data to prevent the mixture of signals originating from the different segments (de Hollander et al., 2015). In all four GLMs (detailed below), task events were convolved with the canonical double-gamma HRF (Glover, 1999), and their temporal derivatives were included. Additionally, the design matrix included temporal derivatives of the main regressors of interest, and (as in the previous analyses) the first five aCompCor components to model physiological noise, seven head motion related parameters, DVARs, and a set of 16 cosines to model low-frequency scanner drifts. The GLMs were fit using ordinary least squares. The estimated BOLD responses, quantified in the GLM’s parameter estimates (betas), were then used as a dependent variable in higher-level analyses to test for functional specialization of STN segments, as detailed below.

##### 2.7.4.1 GLM1: Difficulty and potential payoff cue valence effects

The first GLM included (1) easy and (2) hard stimuli (RDM stimulus onset locked), and (3) payoff and (4) neutral cues (potential payoff cue onset locked) as task events of interest. Firstly, we tested for the presence of the following two contrasts of interest using a within-subject (frequentist and Bayesian) *t*-test for each segment separately: task difficulty (hard – easy), and potential payoff cue type (payoff – neutral). We hypothesized that ‘cognitive’ segment B should respond differently to hard compared to easy trials, and ventromedial ‘limbic’ segment C should show a larger BOLD response after payoff than after neutral cues.

Although these tests indicate whether contrasts were present in each STN segment, the tripartite model of functional specialization holds that the STN segments *differ* in their responses. Suppose that STN segment B indeed has higher BOLD responses for hard than for easy trials, then this would only support functional specialization of segments if the other segments *do not* (or less strongly) show that contrast (Nieuwenhuis et al., 2011). Hence, we then tested for interaction effects between the experimental conditions and the segments using linear MEMs, with task event type, STN segment (A, B or C), and their interaction to predict the BOLD response size:

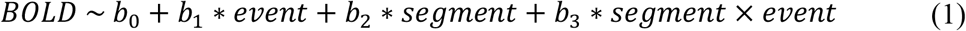

To avoid confusion with the GLM’s *beta* that quantify the BOLD response size, we refer to the MEM parameter estimates with *b*. We fit MEM (1) twice: Once using easy and hard stimuli as events, and once using payoff and neutral cues as predictors. Subjects were included as random intercepts.

Significant contributions of the *segment* × *event* interaction would support functional specialization of segments. We used both frequentist and Bayesian approaches to quantify whether this interaction was supported by the data. Firstly, we estimated the *p*-value associated with the interaction term *segment* × *event*. Since the predictor *segment* consists of three levels, we report *F*-tests using Satterthwaite’s method (Satterthwaite, 1941) to approximate the denominator degrees of freedom. Secondly, we used a Bayesian model comparison approach, in which we sequentially removed individual terms from MEM (1) until the simplest (intercept-only) model remained (i.e., comparing all seven possible models). These models were compared using Bayes factors, which quantify the likelihood ratios of the models (Kass and Raftery, 1995; Ly et al., 2016). Contrary to frequentist *p-*values, Bayes factors can quantify evidence both in favor of and *against* a hypothesis. To describe the strength of evidence (ranging from ‘anecdotal’ to ‘decisive’), we use the conventions proposed by Wetzels and Wagenmakers (2012), who adapted Jeffrey’s (Jeffreys, 1961) original proposal. Bayesian analyses were performed using the BayesFactor (Morey et al., 2018) software for the R programming language (R Core Team, 2017).

##### 2.7.4.2 GLM2: Potential payoff cue lateralization effects

Potential payoff cue-related BOLD effects might be lateralized and represent some a-priori motor facilitation or suppression towards a left or right response. Such a potential payoff cue-related lateralized signal has been shown before in the putamen (Forstmann et al., 2010b) and could potentially cancel out potential payoff cue-related effects when cue direction is ignored. We therefore also fit GLM2 which took into account the direction of the potential payoff cue. It contained the following task regressors: (1) neutral cue, (2) payoff cue (left), (3) payoff cue (right) (all cue onset locked), (4) easy RDM stimulus, and (5) hard RDM stimulus (both RDM stimulus onset locked). The contrast of interest was payoff cue (left) > payoff cue (right).

We tested whether the BOLD responses are different after contralateral potential payoff cues compared to ipsilateral cues, and whether there is any difference therein between STN segments. We constructed a new MEM (1) but using *ipsilateral* and *contralateral* cues (relative to the STN hemisphere) as event types. We used the same frequentist and Bayesian approach as described in 2.7.4.1 to test whether the MEM’s interaction term, which would indicate functional specialization of STN segments, is supported by the data.

##### 2.7.4.3 GLM3: Difficulty effects, de-confounded from error-related processing

As mentioned above, errors are known to cause activity in limbic areas, which could confound drift rate-related activity. To test for activity related to drift rates that cannot be explained by a difference in the error rate, we fit GLM3 that de-confounded the task difficulty from error trials. Here, easy and hard event types were separated by the accuracy of the response on that trial. GLM3 included the following regressors: (1) neutral cue, (2) payoff cue, (3) easy RDM stimulus (correct), (4) easy RDM stimulus (error), (5) hard RDM stimulus (correct), and (6) hard RDM stimulus (error). The two potential payoff cue regressors were locked to the potential payoff cue onset, the four RDM stimulus regressors were locked to the RDM stimulus onset. The main contrast of interest for this GLM was ‘hard RDM stimulus (correct) > easy RDM stimulus (correct)’. To test for the presence of functional subregions, we again constructed a MEM as described equation (1), using the task events ‘hard RDM stimulus (correct)’ and ‘easy RDM stimulus (correct)’. Again, the frequentist and Bayesian methods described in 2.7.4.1 were used to test for functional specialization of STN segments.

##### 2.7.4.4 GLM 4: Motor lateralization effects

To test for lateralized motor activity in the STN, we also fit GLM4, focusing on responses. It included the following regressors: (1) neutral cue, (2) potential payoff cue (both potential payoff cue onset locked), (3) easy trials, (4) hard trials (both RDM stimulus onset locked), (5) left responses, and (6) right responses (both response onset locked). We hypothesized that potential STN ‘motor’ segments should show lateralized motor activity. Specifically, we expected to find increased activity for left versus right responses for right STN segment A versus the left STN segment A, and vice versa. We repeated the LMEM analysis described in 2.7.4.1 but using left response and right response as events of interest. The same frequentist and Bayesian methods described previously were used to test for functional specialization of segments.

#### 2.7.5 Interindividual correlations

We then tested whether interindividual variability in DDM parameters was related to interindividual variability in the corresponding BOLD effects (Forstmann et al., 2011, 2008; Lebreton et al., 2019). The first hypothesis pertained to potential payoff cue-related bias and STN activity. The shift in starting point of every subject was correlated with the contrast ‘payoff cue > neutral cue’ (ROI-wise GLM1). We expected STN ‘limbic’ segment C involved in implementing response biases (the putative ‘limbic’ area) to show such an across-subject correlation between the size of the starting point shift and the size of the BOLD response contrast. We repeated this analysis with the potential payoff cue-related drift rate bias as covariate, with the same hypothesis. Correlations were tested using frequentist and Bayesian tests (Wetzels and Wagenmakers, 2012) as implemented in the BayesFactor software package (Morey et al., 2018).

The second hypothesis pertained to the difficulty manipulation and the resulting drift rate reduction. Specifically, the reduction in drift rate for hard compared to easy trials was correlated, across subjects, with the size of the contrast ‘hard RDM stimulus > easy RDM stimulus’ (ROI-wise GLM1). We expected that activity in the central, ‘cognitive’ STN segment B, should show a significant correlation with the drift rate effect of the difficulty manipulation. We test this hypothesis using the same frequentist and Bayesian correlation tests.

## 3. Results

We first describe three sets of control analyses. Firstly, we test whether the experimental manipulations led to the expected behavioral effects. Secondly, we perform whole-brain voxel-wise GLMs to ensure that the overall patterns of BOLD responses elicited by the experimental manipulations are in line with the literature. Thirdly, we test whether we have sufficient signal quality in the STN segments to be able to detect BOLD responses. After these control analyses, we turn to the confirmatory analysis to test for functional specialization of STN segments.

### 3.1 Behavioral data

The effects of the task manipulations on accuracy and RTs are shown in Figure 2B. We fit a logistic MEM to predict choice accuracy using potential payoff cue congruency and stimulus difficulty. Compared to the model’s intercept (corresponding to the difficult, neutral cue condition), accuracy was higher on easy trials (b = 0.54, SE = 0.06, 95% CI [0.42, 0.66], *p* < 0.001), as well as on trials with a congruent potential payoff cue (b = 0.24, SE = 0.07, 95% CI [0.10, 0.38], *p* < 0.01), but lower on trials with an incongruent potential payoff cue (b = − 0.42, SE = 0.07, 95% CI [-0.56, −0.29], *p* = 0.001). The interaction terms between difficulty and potential payoff cue congruency were not significant.

Secondly, we fit a linear MEM to predict choice RT using potential payoff cue congruency, RDM stimulus difficulty, and choice accuracy as predictors, plus all interactions. The intercept corresponded to correct responses on hard trials with a neutral cue. In this model, we found main effects indicating that RTs were significantly faster for easy trials (b = −0.05, SE = 5.45*10^−3^, 95% CI [-0.06, −0.04], *p* < 0.001) as well as for trials with congruent potential payoff cues (b = −0.02, SE = 6.77*10^−3^, 95% CI [-0.03, −0.01], *p* = 0.004), but slower for errors (b = 0.03, SE = 7.08*10^−3^, 95% CI [0.02, 0.04], *p* < 0.001) and for incongruent trials (b = 0.02, SE = 7.24*10^−03^, 95% CI [0.00, 0.03], *p* = 0.035). Furthermore, we find interactions between choice accuracy and potential payoff cue congruency: In incorrect responses, RTs were *higher* after congruent cues compared to neutral cues (b = 0.03, SE = 0.01, 95% CI [0.01, 0.06], *p* = 0.016) and *lower* after incongruent cues (b = −0.04, SE = 0.01, 95% CI [-0.06, −0.02], *p* < 0.001). Finally, there was an interaction between difficulty and accuracy that indicated that for error responses, the effect of easy trials (versus hard trials) was smaller than for correct responses (b = 0.04, SE = 0.01, 95% CI [0.02, 0.06], *p* < 0.001). The other interaction terms were not significant.

Hence, the effects of the manipulations can be summarized as follows. Hard trials led to less accurate and slower responses, and the effect on RTs was larger for correct compared to error RTs. Furthermore, congruent potential payoff cues increased accuracy, and if an error was made, the corresponding RT was high (compared to neutral cues). Incongruent trials decreased accuracy, and the errors were fast (compared to neutral cues).

The effects of both manipulations are qualitatively consistent with the DDM’s predictions. Choice difficulty is assumed to effect drift rates (e.g., Palmer et al., 2005; Ratcliff and McKoon, 2008; Voss et al., 2004). Increasing the drift rate leads to faster, more accurate responses, which we indeed find in the difficulty manipulation. Potential payoff cues are often thought to cause a bias in the starting point *z* (Diederich and Busemeyer, 2006; Edwards, 1965; Link and Heath, 1975; Mulder et al., 2012; Ratcliff, 1985; Ratcliff and Rouder, 1998; Voss et al., 2004); and/or drift rate (Ashby, 1983; Diederich and Busemeyer, 2006; Mulder et al., 2012; Ratcliff, 1981; Urai et al., 2019). Both biases lead to overall faster responses and more responses for the biased option than for the unbiased option. However, the two types of bias differ in their predictions about the shapes of the RT distributions, as well as about the relative speed of correct and incorrect choices: A start point bias predicts that choices corresponding to the *unbiased* option are slower compared to choices corresponding to the *biased* option, whereas a drift rate bias predicts equally fast biased and unbiased choices.

Hence, formal model fitting was used to discover which type of bias was likely present in the participants’ data. Three DDMs were fit, all assuming that the difficulty manipulation affected drift rates, but assumed that the potential payoff cues biased starting points (DDM1), drift rates (DDM2; i.e., both the potential payoff cue and difficulty manipulation affect drift rates), or both (DDM3). Formal model comparison (Figure 2C) indicated that, when summing BPICs across subjects, the third model provided the most parsimonious trade-off between quality of fit and model complexity. However, a comparison of individual participants’ BPICs (Figure 2C, colors) shows that DDM1 was most parsimonious for 17 participants, DDM2 for 7, and DDM3 for the remaining 9. Thus, there is substantial interindividual variability in the strategies that participants used: A starting point shift was most common, followed by both a combination of starting point and drift rate shifts, but a drift rate shift alone was least common. Most participants (26) used one strategy only, but *across* subjects, DDM3, allowing for both strategies, provided the most parsimonious fit.

To take into account the observed interindividual strategic differences in subsequent analyses, we used Bayesian model averaging (Hoeting et al., 1999) to calculate subject-wise average (across DDMs) parameters. In this approach, a weighted average of each DDM parameter is calculated per subject, with wBPIC-values (Wagenmakers and Farrell, 2004) functioning as weights. Hence, better-fitting DDMs contribute more to the weighted parameter estimate. The averaged drift rate effect of difficulty (*Δv*_*difficulty*_ = *v*_*hard*_ − *v*_*easy*_) and the averaged potential payoff cue-related drift rate (*Δv*_*cue*_) and starting point shift (*Δz*_*cue*_) parameters are shown in Figure 2D.

The posterior predictive distributions in Figure 2E show the quality of fit of DDM3. Whereas DDM3 accounts for the median RTs in most conditions reasonably well, there is a strong misfit in the predicted RT distribution skewness: The DDM predicts much stronger right-skewed distributions than seen in the data, and also misfits the leading edge. Possible causes of this are the (implicit) deadlines of 1.5 seconds and presence of a delay between the choice and feedback, which both have been shown to induce urgency effects that decrease skewness in RT distributions (Evans and Hawkins, 2019; Hawkins et al., 2015; Katsimpokis et al., 2020; Miletić and Van Maanen, 2019). Another indication that an urgency signal influenced decision making are the error RTs, which are slower than the correct RTs. Since urgency entails that the amount of evidence required to inform a decision decreases with decision time, slow decisions are based on less evidence and therefore more likely to be erroneous than faster decisions (Ditterich, 2006; Hawkins et al., 2015).

In summary, the task manipulations led to the expected effects on behavior. While the quality of fit of the DDM could likely be improved by using a model that incorporates urgency, the best-fitting DDM can explain the effects of the manipulations in terms of drift rate and starting point effects.

### 3.2 Whole-brain GLMs

As a second set of control analyses, we fit whole-brain GLMs to test for the effects of the response hand, errors, potential payoff cue, and difficulty. Statistical parametric maps (SPMs) of the main contrasts are shown in Figure 3; a summary of all cluster locations and sizes is presented in Supplementary Table 1. Replicating established findings (Dassonville et al., 1997; Kim et al., 1993a, 1993b), the response hand contrast shows a strong contralateral BOLD response in primary motor (M1) and sensory cortex (S1), as well as in posterior putamen and thalamus. The expected ipsilateral BOLD response is present in the superior cerebellum (Lotze et al., 1999).

An error-related BOLD response was present in insula, anterior cingulate cortex (ACC), and a smaller diffuse cluster in the midbrain. The insula (Eckert et al., 2009; Klein et al., 2013; Ullsperger et al., 2010) and ACC (Botvinick et al., 2001; Duncan and Owen, 2000; Paus, 2001; Paus et al., 1998; Spunt et al., 2012) have frequently been associated with processing of errors and conflicting task demands. A negative BOLD response to errors was present in putamen (which could indicate a negative prediction error (McClure et al., 2003), although these are more commonly associated with ventral striatum), posterior cingulate cortex (speculatively suggesting a lapse of attention on error trials; Leech and Sharp, 2014), and a diffuse cluster in left occipital cortex. The latter cluster might be the result of attention-lapse related activity modulations of the visual system (Reynolds and Heeger, 2009) and/or involuntary saccades that impaired task performance (Sylvester et al., 2005).

Payoff cues caused an increased BOLD response compared to neutral cues in the superior parietal cortex, left premotor cortex, and pre-SMA. Parietal cortex has often been implicated in evidence accumulation during perceptual decision making (Gold and Shadlen, 2001; Kim and Shadlen, 1999; O’Connell et al., 2012; Shadlen and Newsome, 2001), and the increased BOLD response could reflect an increased starting point or drift rate. However, the model-based analysis showed no evidence that the BOLD response contrast size covaried with either the size of the shifts in starting point or drift rate. The increased pre-SMA and premotor BOLD responses likely reflect an increased motor readiness in response to a payoff cue (Cunnington et al., 2005, 2002, 1996; Pedersen et al., 1998). Contrary to expectations, we did not find potential payoff cue-related BOLD responses in limbic areas such as the orbitofrontal cortex (c.f. Forstmann et al., 2010b; Keuken et al., 2014b; Summerfield and Koechlin, 2010).

The difficulty manipulation caused an increased BOLD response in hard trials compared to easy trials in anterior insula (over and above the effect of errors), as commonly found using difficulty manipulations with a variety of decision-making tasks (Keuken et al., 2014b). Furthermore, the model-based analysis shows that in right insula, the size of the BOLD response difference (hard – easy) covaries with the drift rate difference (*v*_*hard*_ − *v*_*easy*_), indicating that participants with a larger difference in drift rates between conditions, also show a larger difference in right insular BOLD responses. The right insula specifically has earlier been implicated in drift rate effects in perceptual decision making (Ho et al., 2009). No evidence was found for difficulty-related BOLD responses in the dorsolateral prefrontal cortex (Heekeren et al., 2004).

In sum, the whole-brain results partly led to the expected effects of the task manipulations, consistent with the literature. Unfortunately, no difficulty-related BOLD responses in the dorsolateral prefrontal cortex were found, and the model-based analysis of the effect of the potential payoff cue showed no regions in which BOLD response size covaried with the behavioral effect size. Note that the payoff cue manipulation is known to be relatively weak compared to cue-based probability manipulations (Forstmann et al., 2010b; Mulder et al., 2012; Voss et al., 2004). Here, we preferred the payoff manipulation to activate limbic processing. Furthermore, the two different strategies of incorporating the payoff information in decision-making behavior (via a starting point or drift rate bias) may rely on partially different brain networks, which could weaken statistical power to find a covariances with DDM parameters. In the supplementary materials, we report SPMs of the limbic manipulation with a liberal, exploratory threshold.

### 3.3 Deconvolution analyses

As a third control analysis, we aimed to test whether there was sufficient BOLD sensitivity in the STN, and whether the shape of the BOLD response was in line with the canonical, double-gamma HRF typically used in GLM analyses. To do so, we applied a deconvolution analysis to the mean timeseries of each STN segment. The fit deconvolution model was visualized by plotting the percentage signal change as a function of time after onset of the event, as shown in Figure 4. All STN segments show a BOLD response to the RDM stimulus. Furthermore, the shape of the BOLD responses was in line with the canonical double-gamma BOLD response, although we do note between-region variability in the shape. Specifically, in left segments A and B, and in right segment C, the size of the post-stimulus undershoot was larger than commonly assumed in the canonical HRF (de Hollander et al., 2017; Miletić et al., 2020). Between-region variability in the shape of the HRF has been found before (Boillat and Van der Zwaag, 2019; Gonzalez-Castillo et al., 2012; Handwerker et al., 2004; Miezin et al., 2000). Furthermore, it appears that the peak of the BOLD response occurs roughly 4 seconds after RDM stimulus onset, thus earlier than the canonical 6 seconds. However, some caution must be taken in interpreting this difference, since the TR of 3 seconds is relatively high, which may limit our ability to determine the exact time at which the response peaks. Contrary to the RDM stimulus, the potential payoff cue seems to evoke no typical BOLD response in most segments (in line with Keuken et al., 2015), except for a potential trend towards a BOLD response in left STN ventromedial segment A.

**Figure 4.**
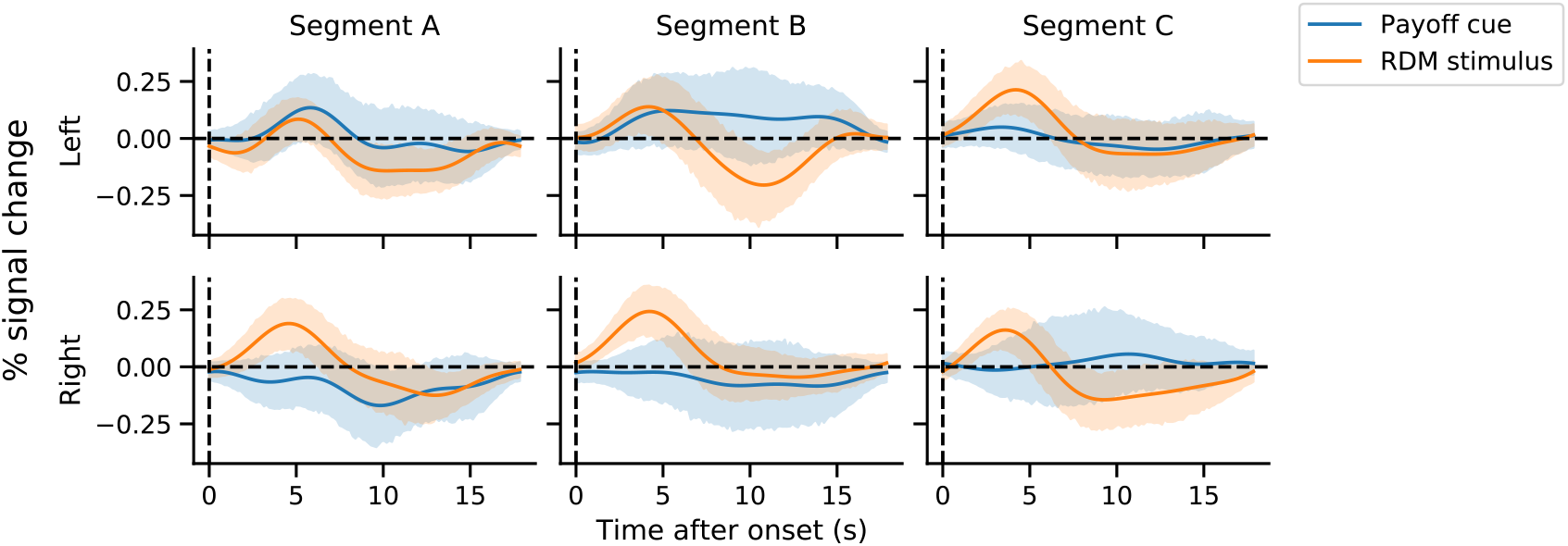
Deconvolution of BOLD-responses to the potential payoff cue (blue) and RDM stimulus (orange) using Fourier basis sets. Shaded areas correspond to 95% confidence intervals.

### 3.4 ROI-wise GLMs

Having confirmed that fMRI signal quality was sufficient to detect BOLD responses in the individual segments of the STN, we fit four GLMs to test our main hypotheses. Figure 5 summarizes all four GLMs’ BOLD response estimates. These BOLD response estimates were used as dependent variables. In Table 1, we present per-segment, within-subject *t*-tests for all contrasts of interest. Using linear MEMs, we then tested for differences between conditions and differences between segments.

**Table 1.**
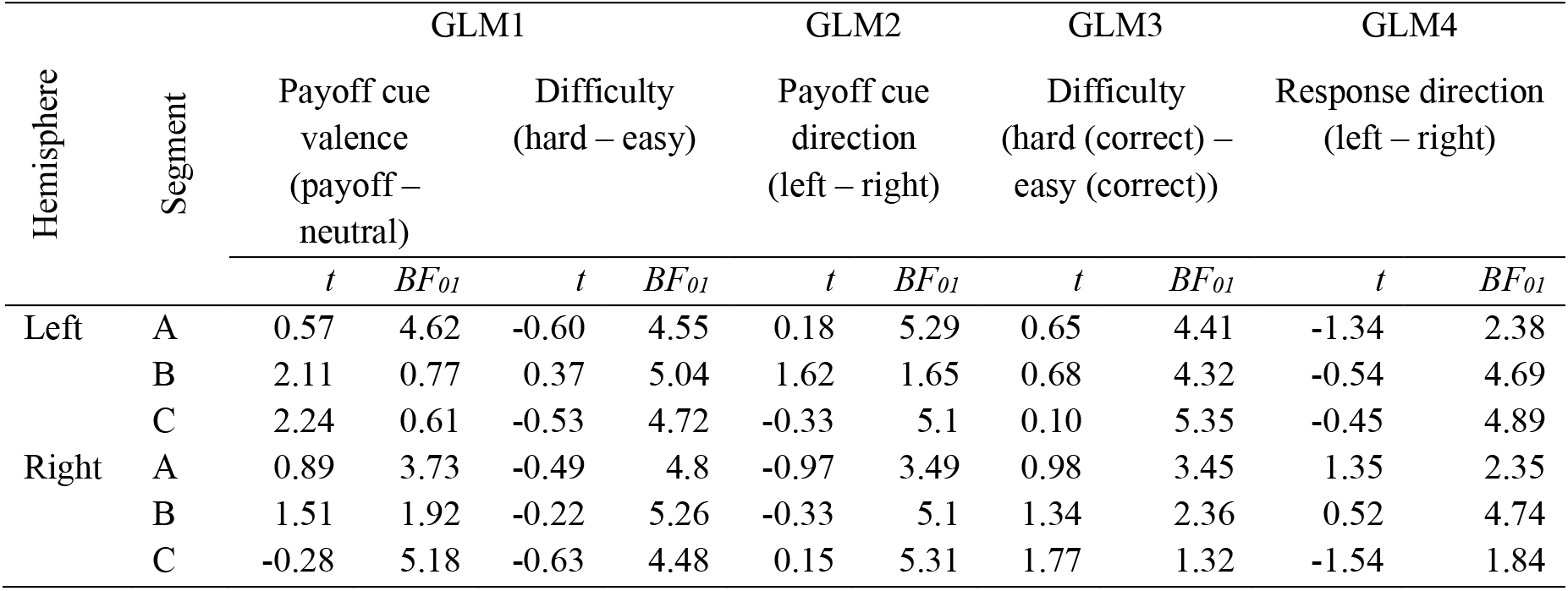
Results of per-subregion within-subject frequentist and Bayesian *t*-tests. Frequentist *t*-tests had 32 degrees of freedom, and none were significant after correcting for the false discovery rate (*q* < 0.05). Bayes factors are presented as BF_01_, where values greater than 1 indicate evidence against the presence of a difference, and values lower than 1 evidence in favor of a difference.

| Hemisphere | Segment | GLM1 |  |  |  | GLM2 |  | GLM3 |  | GLM4 |  |
| --- | --- | --- | --- | --- | --- | --- | --- | --- | --- | --- | --- |
|  |  | Payoff cue valence (payoff – neutral) |  | Difficulty (hard – easy) |  | Payoff cue direction (left – right) |  | Difficulty (hard (correct) – easy (correct)) |  | Response direction (left – right) |  |
|  |  | *t* | *BF*<sub>01</sub> | *t* | *BF*<sub>01</sub> | *t* | *BF*<sub>01</sub> | *t* | *BF*<sub>01</sub> | *t* | *BF*<sub>01</sub> |
| Left | A | 0.57 | 4.62 | -0.60 | 4.55 | 0.18 | 5.29 | 0.65 | 4.41 | -1.34 | 2.38 |
|  | B | 2.11 | 0.77 | 0.37 | 5.04 | 1.62 | 1.65 | 0.68 | 4.32 | -0.54 | 4.69 |
|  | C | 2.24 | 0.61 | -0.53 | 4.72 | -0.33 | 5.1 | 0.10 | 5.35 | -0.45 | 4.89 |
| Right | A | 0.89 | 3.73 | -0.49 | 4.8 | -0.97 | 3.49 | 0.98 | 3.45 | 1.35 | 2.35 |
|  | B | 1.51 | 1.92 | -0.22 | 5.26 | -0.33 | 5.1 | 1.34 | 2.36 | 0.52 | 4.74 |
|  | C | -0.28 | 5.18 | -0.63 | 4.48 | 0.15 | 5.31 | 1.77 | 1.32 | -1.54 | 1.84 |

**Figure 5.**
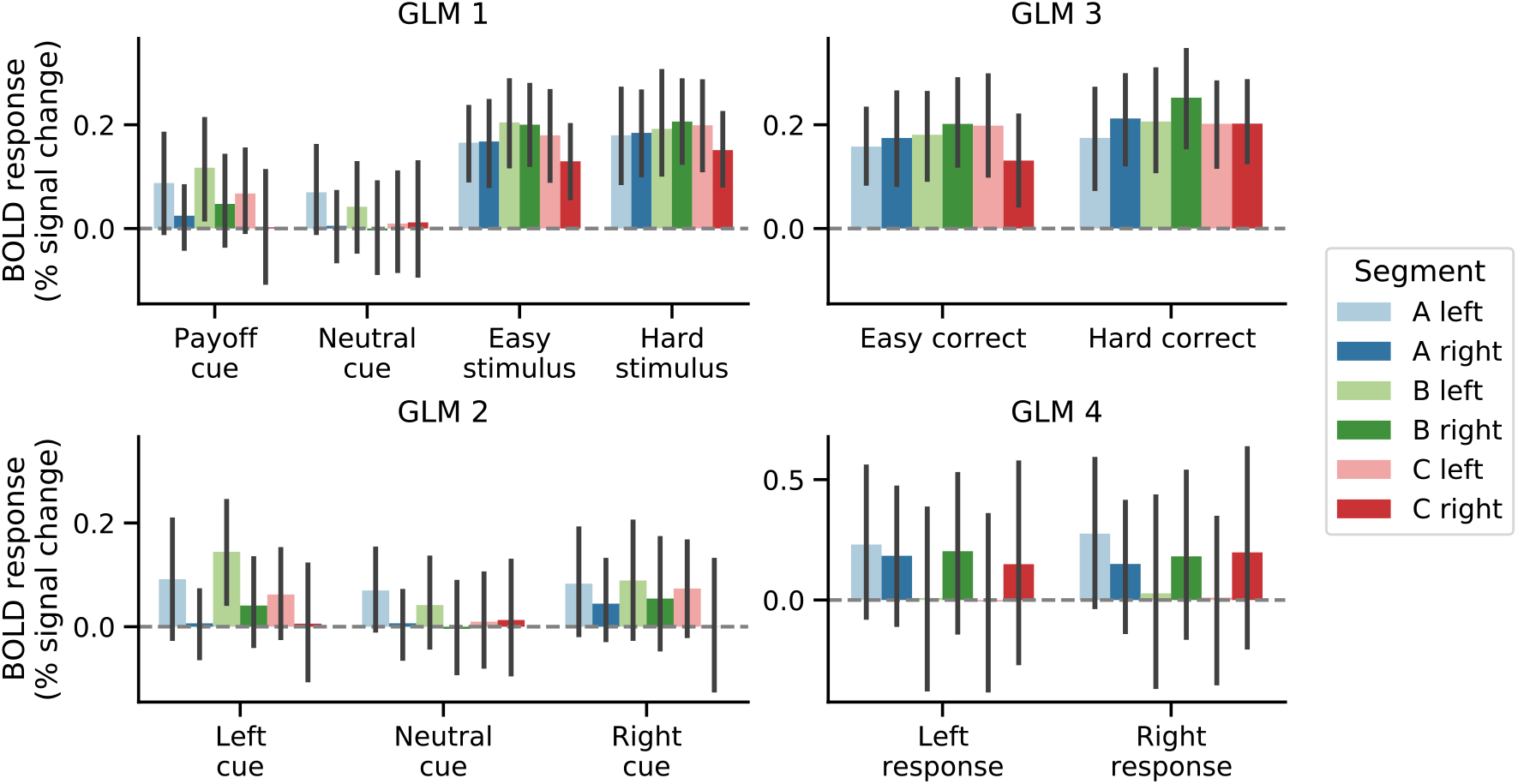
Parameter estimates from four GLMs fit to the timeseries of each subregion within the STN. Error bars indicate 95% confidence intervals.

First, we used GLM1 to test whether potential payoff cue valence affected BOLD responses, and whether there were between-segment differences in these BOLD responses. A linear MEM showed no evidence for an effect of potential payoff cue type (payoff vs. neutral; *F*(1, 160) = 3.49, *p* = 0.064), of segment (A, B or C; *F*(2, 160) = 0.85, *p* = 0.428), nor, critically, of their interaction (*F*(2, 160) = 0.54, *p* = 0.58) on the BOLD response size. Bayesian model comparison furthermore preferred an intercept-only model over all seven possible alternative models that included potential payoff cue type, STN segment, and/or their interaction predictors. Specifically, there was substantial evidence against a model with potential payoff cue type as predictor (BF_01_ = 3.605), strong evidence against a model with STN segment as predictor (BF_01_ = 14.74), substantial evidence against a model with their interaction as predictor (BF_01_ = 9.277), and even stronger evidence against more complex MEMs (all BF_01_ > 34). Hence, the BOLD responses to payoff and neutral cues were the same in all STN segments.

Second, we used GLM1 to test whether RDM stimulus difficulty affected STN BOLD responses, using the same analysis but including easy and hard stimuli as event types. A linear mixed effects model also indicated no evidence for an effect of difficulty (*F*(1, 160) = 0.3, *p* = 0.586), nor of STN segment (*F*(2, 160) = 1.157, *p* = 0.32), or their interaction (*F*(2, 160) = 0.13, *p* = 0.88) on BOLD response size. Bayesian model comparison again favored an intercept-only model. There was substantial evidence against an influence of difficulty (BF_01_ = 5.96), strong evidence against a difference between segments (BF_01_ = 10.59), and against their interaction (BF_01_ = 10.19). There was even stronger evidence against the more complex models including multiple predictors (all BF_01_ > 62). Thus, the BOLD responses to hard and easy stimuli were the same in all STN segments.

Third, we used GLM2 to test for an effect of the potential payoff cue direction on BOLD response sizes. Here, a linear MEM predicting the BOLD response suggested no effect of potential payoff cue laterality (ipsilateral vs contralateral relative to the STN hemisphere; *F*(1, 160) = 0.64, *p* = 0.436), no effect of STN segment (*F*(2, 160) = 1.75, *p* = 0.18), nor an interaction (*F*(2, 160) = 0.40, *p* = 0.67). Bayesian model comparison again preferred an intercept-only model, with substantial evidence against a model including potential payoff cue laterality as predictor (BF_01_ = 5.80), strong evidence against a model including STN segment (BF_01_ = 10.94), substantial evidence against a model with an interaction between subregion and laterality (BF_01_ = 9.63), and stronger evidence against all more complex models (all BF_01_ > 54). Thus, we also found that all STN segments responded equally to contralateral and ipsilateral potential payoff cues.

Fourth, we used GLM3 to test whether difficulty affects the STN when only the trials are considered in which the participants gave correct responses, to de-confound the effect of error-related processing. Hence, the MEM with difficulty (hard, correct vs easy, correct), STN segments, and their interaction as predictors, showed no effect of difficulty (*F*(1, 160) = 2.69, *p* = 0.1), of STN segment (*F*(2, 160) = 0.846, *p* = 0.43), nor their interaction *(F*(2, 160) = 0.03, *p* = 0.97). Bayesian model comparison favored a null model again, with substantial evidence against an effect of difficulty (BF_01_ = 3.21), strong evidence against an effect of STN segment (BF_01_ = 12.76), strong evidence against an interaction effect (BF_01_ = 10.69), and strong to decisive evidence against all more complex models including multiple predictors (all BF_01_ > 35). Thus, also when considering only the trials with correct responses, all STN segments responded equally to easy and hard trials.

Fifth, we tested for motor lateralization effects using GLM4. Here, we constructed a linear mixed effects model predicting the BOLD response size using the response direction (contralateral vs ipsilateral) and subregion. Again, we found no main effects of response direction (*F*(1, 358) = 0.04, *p* = 0.85), no main effect of subregion (*F*(2, 358) = 1, *p* = 0.37), nor an interaction (*F*(2, 358) = 0.05, *p* = 0.95). Bayesian model comparisons favored a null model again, with substantial evidence against a model including a main effect of response direction (BF_01_ = 8.91), strong evidence against a model including a main effect of subregion (BF_01_ = 21.8), strong evidence against a model including their interaction (BF_01_ = 18.8), and decisive evidence against more complex models including multiple predictors (all BF_01_ > 175).

To summarize, in this section, frequentist tests suggested no evidence for effects of any of our manipulations (difficulty, potential payoff cue valence, motor response direction) on the STN BOLD responses, and no evidence for differences between the segments or for interactions. The Bayesian analyses consistently showed substantial to strong evidence against such effects.

### 3.5 Interindividual differences

Next, we took an interindividual differences approach to analyze the data, reasoning that the size of the behavioral differences should covary with the size of the BOLD response differences in the STN. Specifically, we tested whether the size of the shift in the DDM’s starting points and drift rates due to a potential payoff cue was correlated to the difference in BOLD responses between payoff and neutral cues (obtained from GLM1). Furthermore, we tested whether the difference between the DDM’s drift rates (hard – easy) was correlated with the corresponding difference in BOLD responses (also obtained from GLM1). All correlations, including their coefficients and corresponding Bayes factors, are shown in Figure 6. Frequentist tests showed that none of the correlation coefficients was significant, and the Bayes factors indicated anecdotal evidence against correlations. In summary, there was no evidence for any interindividual correlations between the size of the behavioral effect and the size of the STN’s BOLD responses in all subregions.

**Figure 6.**
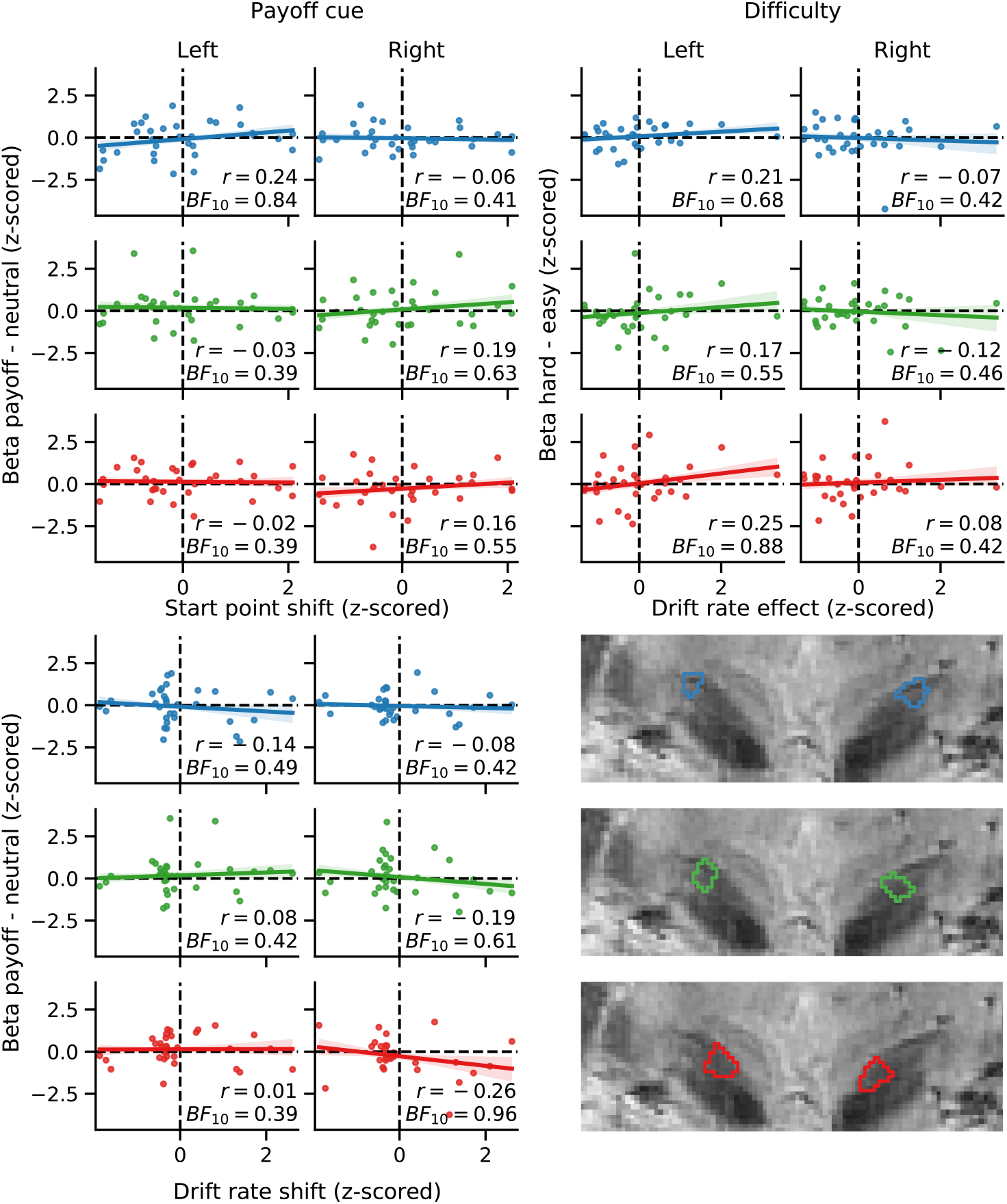
Interindividual correlations between BOLD-response contrasts and parameter estimate contrasts. Both the BOLD-response contrasts (y-axis) and the parameter estimate contrasts (x-axis) are *z*-scored. Colors indicate subregions A (blue), B (green), and C (red). Shaded areas correspond to 67% confidence intervals. Pearson’s correlation coefficient *r* is included in each panel, with the corresponding Bayes factors.

## 4 Discussion

A prominent hypothesis on the organization of the STN holds that it contains three functionally distinct subregions, respectively specialized in cognitive, limbic, and motor processing (Temel et al., 2005a). Here, we tested this hypothesis in healthy volunteers using ultra-high field 7 T BOLD-fMRI with protocols optimized to study anatomy and function of STN (de Hollander et al., 2017, 2015; Miletić et al., 2020; Mulder et al., 2019), manual delineations to maximize anatomical accuracy, and a well-understood task with manipulations hypothesized to elicit cognitive, limbic, and motor effects on behavior.

Specifically, we used a random dot motion decision-making task with a payoff bias and a difficulty manipulation, and we furthermore analyzed the different response directions. Control analyses confirmed that the task manipulations elicited the expected behavioral effects. Furthermore, whole-brain analyses of the functional neuroimaging data directly replicated well-established findings on the effect of motor lateralization. The difficulty manipulation caused differential BOLD responses in right anterior insula, and the size of the BOLD response difference covaried with the size of the behavioral difficulty effect. Drift rate effects on anterior insula have frequently been reported in decision-making tasks with difficulty manipulations (Keuken et al., 2014b). The payoff cues resulted in increased BOLD responses in parietal cortex, which is commonly implicated in evidence accumulation in perceptual decision making, but the presence of between-subject strategy differences limited our ability to directly compare our results with the (relatively scarce) literature on this manipulation. Deconvolution analyses confirmed that we were able to detect BOLD responses in all STN segments. Despite the promising results of our control analyses, we found no evidence for functionally specialized subregions in the STN. More strongly, Bayesian analyses repeatedly suggested substantial to strong evidence against functionally distinct subregions in the STN.

Our results seem contradictory with studies that suggests the presence of subregions (for recent overviews, Emmi et al., 2020; van Wijk et al., 2020). The tripartite model was developed partly based on anatomical anterograde tracing studies (Haynes and Haber, 2013; Joel and Weiner, 1997; Parent and Hazrati, 1995), which showed that the termination sites in the STN from various cortical regions are organized along the STN’s dorsolateral-ventromedial axis. ‘Motor’ cortical areas (M1, SMA) predominantly terminate in the dorsolateral part, ‘limbic’ cortical areas (e.g., vmPFC/OFC) terminate in the ventromedial part, and ‘cognitive’ areas (ACC and dPFC) in between, although the projections are highly overlapping (Alkemade, 2013; Haynes and Haber, 2013; Lambert et al., 2015).

Other studies that are considered to support a three-partite functional organization include LFP recordings from DBS electrodes in PD patients. Various studies report the peak amplitude of beta oscillations are near the dorsolateral border (de Solages et al., 2011; Kühn et al., 2005; Seifried et al., 2012; Trottenberg et al., 2007; Weinberger et al., 2006; Zaidel et al., 2010), and alpha peak amplitude more towards the ventromedial border (Horn et al., 2017). Yet, there is substantial variability in peak site locations, and because DBS surgery targets the dorsolateral part of the STN, the ventromedial tip is inherently undersampled, complicating the interpretation of these findings in terms of the functional organization of the STN (van Wijk et al., 2020).

It is possible that the STN is indeed specialized in functionally distinct subregions, but that we were unable to find these due to experimental design choices. One important consideration is how we operationalized the hypothesized subregions of the STN into three segments. Here, we used a PCA on the individual STN masks to identify the ventromedial-dorsolateral axis, along we divided the STNs in three segments of equal volume. Some other studies suggest more complex shapes of subregions (e.g., Ewert et al., 2018; Horn et al., 2017; but see Lambert et al., 2012), but there is considerable variability in the reported shapes and even number of hypothesized subdivisions (for review, see Keuken et al., 2012). Other work suggests the STN is organized along a gradient of changes with no clear boundaries (Alkemade et al., 2019; de Hollander et al., 2014). Our approach provides a principled way to formalize the hypothesis that the ventromedial-dorsolateral axis is the most important axis of organization. An alternative analysis strategy could have been to use purely voxel-based analyses and define three subdivision ROIs in a more data-driven manner. However, preliminary analyses clearly showed that the signal-to-noise ratio of the BOLD signal at a field strength of 7 Tesla is not reliable enough to elucidate activation patterns in single voxels within single individuals. Therefore, to improve experimental power and to refrain from overly exploratory analysis approaches, we opted to use the ROI-based approach employed here and advocated elsewhere (see also de Hollander et al., 2017, 2015).

A second design consideration is our operationalization of the cognitive, limbic, and motor domains in the experimental paradigm. We used the random-dot motion task since the STN is often thought to play a central role in speeded (perceptual) decision making (Bogacz, 2007; Bogacz and Gurney, 2007; Frank, 2006; Frank et al., 2015; Green et al., 2013; Keuken et al., 2015, 2018b; Mink, 1996). The operationalizations of the cognitive domain by means of a difficulty manipulation, of the limbic domain by means of the potential reward, and the motor domain using the response hands, were based on earlier studies using similar manipulations (Mulder et al., 2014). The difficulty manipulation has previously been shown to affect ‘cognitive’ brain areas such as the dorsolateral prefrontal cortex (Heekeren et al., 2004; Kaiser et al., 2007). Earlier work also suggested that STN activity reflects a ‘conflict’ or ‘normalization’ signal (e.g., Bogacz and Gurney, 2007; Frank et al., 2015; Keuken et al., 2015) that should be inversely proportional to choice difficulty. The detection of conflicts allows the STN to increase response thresholds (e.g. Bogacz and Gurney, 2007; Frank et al., 2015; Herz et al., 2016) to prevent premature, inappropriate responses. This proposed inhibitory role of the STN extends to other cognitive tasks (Soh and Wessel, 2021; Wessel et al., 2016), and could be facilitated by the short neural pathways between the visual system and the STN (Coizet et al., 2009) which could allow the STN to quickly detect whether a stimulus is sufficiently clear to allow for a fast response, or whether thresholds should be increased.

The potential reward manipulation has been shown to affect ‘limbic’ brain areas such as the orbitofrontal cortex (Forstmann et al., 2010a; Summerfield and Koechlin, 2010). Unfortunately, we were not able to replicate these findings. We found significant task-related BOLD responses in the STN, but found no differences in the BOLD response between hard and easy trials, between payoff and neutral cues, or between left and right responses. It is possible that the tripartite model is correct, but that our manipulations (despite being tailored to do so) failed to tap into the three distinct cortico-basal ganglia networks, hence failing to elicit specific cognitive, limbic, and motor functioning of the STN. If so, it is paramount to augment the tripartite model with specific attributions of cognitive, limbic, and motor functions, such that the model remains testable. Then, future work can assess whether other experimental designs can elicit BOLD responses that support a tripartite model.

Our study is the first to test for functional specialization of STN subregions using ultra-high field 7 T BOLD-fMRI, which, contrary to LFP recordings, allowed for testing healthy volunteers and sample across the entire STN. However, the spatial resolution of the BOLD response can be lower than the nominal scanning resolution due to draining veins that cause BOLD responses in regions downstream from the activated neural tissue (Turner, 2002; Uǧurbil et al., 2003). One study at 7 T estimates the full-width-half-maximum of the BOLD point-spread function in cortical gray matter to be in the range of 1.5 – 2 mm (Shmuel et al., 2007). This is a known issue for laminar fMRI, where non-BOLD fMRI methods (Huber et al., 2019, 2018, 2015, 2014) are more frequently used to disentangle neural activity from the different cortical layers using sub-millimeter scanning protocols. However, the estimated point spread function is still smaller than the length of the STN along the ventromedial-dorsolateral axis (approximately 1 cm). Another consideration here is the use of partial Fourier (required to obtain a TE of 14 ms), which decreases the effective resolution in the phase encoding direction (anterior-posterior) in our functional data. The STN, however, extends across multiple slices (see Figure 1), which increases the number of independent samples from the different segments. Furthermore, compared to lower field strengths, 7 T fMRI appears to be less sensitive to macrovasculature, since the T_2_* of macrovasculature is relatively much lower compared to the microvasculature at 7 T (Markuerkiaga et al., 2016; Uludaǧ et al., 2009). In future work, it will be of utmost importance to investigate the neurovascular coupling in and surrounding the STN in order to understand the limitations of the spatial specificity of the BOLD response in such subcortical regions.

Another limitation of using BOLD-fMRI to study the STN is the gradient of increasing iron concentrations towards the ventromedial tip of the structure (de Hollander et al., 2014; Stüber et al., 2014). The presence of iron in brain tissue decreases the transversal relaxation time T_2_*, which in turn affects the size of the BOLD response (under a constant echo time): Under the assumption of mono-exponential signal decay, signal-to-noise ratios decrease exponentially with echo time, but BOLD contrast increases linearly (e.g., Kundu et al., 2012). Hence, voxels with relatively low T_2_* values are expected to have a relatively large BOLD response (e.g., in percent signal change) and relatively large amount of signal noise. Optimal BOLD contrast-to-noise ratios (and therefore, highest first-level *t*-values) are reached when the echo time is equal to the T_2_* of the voxel (Posse et al., 1999). Therefore, it can be expected that both the BOLD contrast estimates as well as the *t*-values vary along the ventromedial-dorsolateral axis of the STN. Future studies supporting the tripartite model using BOLD-fMRI should therefore ensure that the T_2_* variability within the STN did not influence the results.

In sum, we tested for the presence of three functional subdivisions in the STN using ultra-high field 7 T BOLD-fMRI in healthy volunteers with a task paradigm. The results did not support functional subdivisions, and more generally show no BOLD responses in the STN related to task difficulty, payoff cues, or motor response direction. It is important for future work that the tripartite model is further augmented to clarify which experimental paradigms and manipulations are expected to lead to differential activity between subregions, such that these predictions can be tested.

## Acknowledgements

We would like to thank Domenica Wilfing, Elisabeth Wladimirow, and Josephine Groot for their help in acquiring the data. We like to thank Wietske van der Zwaag for helping to implement the functional data acquisition protocol for dataset 2. This research was supported by an ERC-stg (313481; BUF) and NWO-VICI grant (016.Vici.185.052; BUF) and an NWO-Rubicon grant (GdH).

## Code & Data availability statement

All code used to analyze the data can be found on https://osf.io/nurde/. Due to restrictions from GDPR, we are unable to make dataset 1 publicly available. Dataset 2 can be found on https://osf.io/nurde/.

## Supplementary Materials

**Figure S1.**
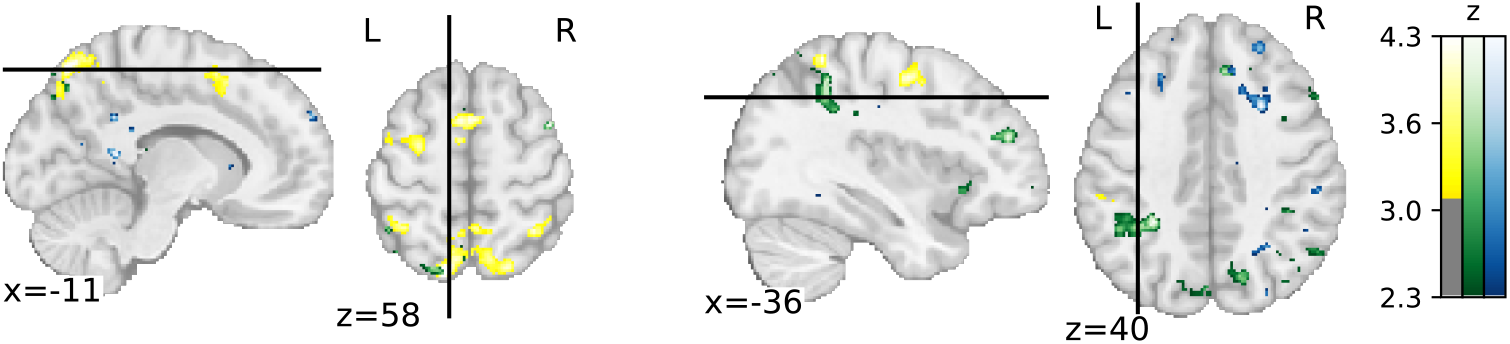
SPMs of the *payoff cue > neutral cue* contrast, against the group-level intercept (yellow), and covariance with the corresponding subject-wise drift rate shifts (green) and starting point shifts (blue). The SPM of the group-level intercept was (as in the main text) corrected for the family-wise error rate by first thresholding at the voxel level (z > 3.1, corresponding to *p* < 0.001) and then at the cluster level (*p* < 0.05, two-sided). The other SPMs were liberally thresholded at z > 2.3 (corresponding to *p* < 0.05) and no correction for multiple comparisons was applied. These SPMs suggest that (without correction for multiple comparisons) there was a covariance between the potential payoff cue-related BOLD and drift rate differences in parietal cortex, partially overlapping with regions that showed a main effect of the *payoff > neutral* cue contrast (in yellow). Without correction for multiple comparisons, the largest cluster of voxels (259 voxels) that would suggest covariance between potential payoff cue-related BOLD and starting point shifts, was centered on white matter in the right frontal lobe (rightmost panel). The other clusters are small ( < 150 voxels = 506.25 mm^3^) and scattered across the brain. Combined, this suggests that even with a liberal threshold, no clusters would be found that support a covariance between BOLD response differences (*payoff > neutral* cues) and the corresponding starting point shifts.

**Supplementary Table S1.** All significant clusters from the four whole-brain GLM contrasts (Figure 3 in the main text). X, Y, Z indicate the coordinates of the voxel with the peak BOLD contrast in MNI2009c-space (1 mm). Maximum or minimum z-values are reported for positive and negative contrasts, respectively.

| | X | Y | Z | Volume ($\text{mm}^3$) | Max / min z-value |
| --- | --- | --- | --- | --- | --- |
| *Left > right response* |  |  |  |  |  |
| Left Pre/Postcentral Gyrus (M1/S1) | -40 | -22 | 55 | 32005.125 | -7.83 |
| Right Pre-/Postcentral gyrus (M1/S1) | 46 | -32 | 61 | 25525.125 | 7.20 |
| Right Insula | 35 | -16 | 17 | 11981.25 | 6.30 |
| Left cerebellum | -22 | -53 | -20 | 10438.875 | 8.37 |
| Right cerebellum | 25 | -49 | -28 | 9784.125 | -7.47 |
| Left Central Opercular Cortex | -38 | -20 | 22 | 6780.375 | -6.04 |
| Right Cingulate Gyrus | 5 | -2 | 46 | 3776.625 | 5.44 |
| Left Precentral Gyrus | -7 | -26 | 47 | 3182.625 | -6.42 |
| Right thalamus | 16 | -20 | 4 | 3159 | 7.32 |
| Left thalamus | -19 | -20 | 8 | 1748.25 | -6.27 |
| Right white matter | 20 | -22 | 31 | 1194.75 | -4.98 |
| Right Lateral Occipital Cortex | 22 | -76 | 31 | 1140.75 | -4.22 |
| Left Occipital Pole | -7 | -98 | 13 | 1096.875 | -4.35 |
| Right Precentral Gyrus | 10 | -26 | 50 | 1066.5 | 6.23 |
| Right Occipital Pole | 19 | -94 | 2 | 688.5 | -4.41 |
| *Error* |  |  |  |  |  |
| Posterior Cingulate Gyrus | 1 | -41 | 32 | 20604.375 | -6.80 |
| Paracingulate Gyrus / Pre-SMA | 11 | 23 | 35 | 16281 | 6.99 |
| Right inferior frontal gyrus | 32 | 17 | 13 | 14512.5 | 7.25 |
| Left inferior parietal lobule | -40 | -73 | 47 | 11711.25 | -6.19 |
| Right insula | -32 | 28 | 2 | 9834.75 | 7.83 |
| Left Inferior Temporal Gyrus<br>(temporooccipital) | -53 | -55 | -22 | 8386.875 | -5.75 |
| Brainstem / Thalamus | -8 | -29 | -11 | 8211.375 | 5.72 |
| Left Middle Frontal Gyrus | -34 | 16 | 59 | 8002.125 | -5.83 |
| Left putamen | -31 | -16 | -1 | 6692.625 | -5.92 |
| Right putamen | 28 | -10 | 4 | 6010.875 | -5.74 |
| Left Frontal Pole | -10 | 65 | 14 | 3901.5 | -4.91 |
| Right Superior Frontal Gyrus | 20 | 31 | 50 | 3429 | -5.20 |
| Left frontal orbital cortex | -31 | 32 | -16 | 2446.875 | -5.02 |
| Right Posterior Supramarginal Gyrus | 64 | -43 | 28 | 2187 | 4.60 |
| Left Inferior Frontal Gyrus | -56 | 32 | 14 | 2068.875 | -5.63 |
| Right Middle Frontal Gyrus | 44 | 17 | 29 | 2021.625 | 5.45 |
| Left Anterior Parahippocampal Gyrus | -20 | -13 | -26 | 1856.25 | -4.97 |
| Right Anterior Parahippocampal Gyrus | 20 | -11 | -26 | 1741.5 | -5.15 |
| Left Intracalcarine Cortex | -16 | -76 | 13 | 1653.75 | 4.75 |
| Right Planum Temporale | 64 | -14 | 8 | 1238.625 | -4.75 |
| Right Intracalcarine Cortex | 8 | -67 | 11 | 1103.625 | 4.92 |
| Left Occipital cortex (V4/V2) | -37 | -92 | -14 | 1053 | -4.37 |
| Right Middle Temporal Gyrus<br>(temporooccipital) | 67 | -46 | -8 | 644.625 | -4.89 |
| Right Frontal Pole | 34 | 38 | -10 | 607.5 | -4.52 |
| Right Posterior Middle Temporal Gyrus | 52 | -23 | -8 | 550.125 | 4.28 |
| Left Posterior Parahippocampal Gyrus | -32 | -38 | -13 | 523.125 | -4.47 |
| Right inferior parietal lobule | 49 | -65 | 46 | 378 | -4.09 |
| *Payoff > Neutral cue* |  |  |  |  |  |
| Superior parietal lobule (bilateral) | -16 | -76 | 58 | 5113.125 | 5.30 |
| Left Precentral Gyrus | -43 | -5 | 55 | 3256.875 | 4.52 |
| Pre-supplementary motor area | -5 | 8 | 56 | 2281.5 | 4.40 |
| Lateral Occipital Cortex, inferior division<br>(V5) | -50 | -73 | -2 | 1164.375 | 5.10 |
| Superior Parietal Lobule / Anterior intra-<br>parietal sulcus | -37 | -50 | 59 | 840.375 | 4.30 |
| Right inferior frontal gyrus | 56 | 28 | -2 | 580.5 | -4.55 |
| *Hard > easy RDM stimulus* |  |  |  |  |  |
| Left insula | -37 | 19 | -2 | 1218.375 | 4.55 |
| Right insula | 31 | 23 | -2 | 1046.25 | 4.54 |
| Right Lingual Gyrus | 19 | -73 | -7 | 772.875 | -4.39 |
| Lateral Superior Occipital Cortex | 20 | -88 | 26 | 675 | -4.54 |
| *Hard > easy RDM stimulus (drift rate covariance)* |  |  |  |  |  |
| Right Insula | 35 | 22 | -5 | 1134 | 4.21 |

## References

Abraham, A., Pedregosa, F., Eickenberg, M., Gervais, P., Mueller, A., Kossaifi, J., Gramfort, A., Thirion, B., Varoquaux, G., 2014. Machine learning for neuroimaging with scikit-learn. Front. Neuroinform. 8, 14. https://doi.org/10.3389/fninf.2014.00014

Alexander, G.E., Crutcher, M.D., 1990. Functional architecture of basal ganglia circuits: neural substrates of parallel processing. Trends Neurosci. 13, 266–271. https://doi.org/10.1016/0166-2236(90)90107-L

Alexander, W.H., Brown, J.W., 2011. Medial prefrontal cortex as an action-outcome predictor. Nat. Neurosci. 14, 1338–1344. https://doi.org/10.1038/nn.2921

Alkemade, A., 2013. Subdivisions and anatomical boundaries of the subthalamic nucleus. J. Neurosci. 33, 9233–9234. https://doi.org/10.1523/JNEUROSCI.1266-13.2013

Alkemade, A., de Hollander, G., Miletic, S., Keuken, M.C., Balesar, R., de Boer, O., Swaab, D.F., Forstmann, B.U., 2019. The functional microscopic neuroanatomy of the human subthalamic nucleus. Brain Struct. Funct. 224, 3213–3227. https://doi.org/10.1007/s00429-019-01960-3

Alkemade, A., Mulder, M.J., Groot, J.M., Isaacs, B.R., van Berendonk, N., Lute, N., Isherwood, S.J., Bazin, P.-L., Forstmann, B.U., 2020. The Amsterdam Ultra-high field adult lifespan database (AHEAD): A freely available multimodal 7 Tesla submillimeter magnetic resonance imaging database. Neuroimage 221, 117200. https://doi.org/10.1016/j.neuroimage.2020.117200

Ando, T., 2007. Bayesian predictive information criterion for the evaluation of hierarchical Bayesian and empirical Bayes models. Biometrika 94, 443–458. https://doi.org/10.1093/biomet/asm017

Aquino, D., Bizzi, A., Grisoli, M., Garavaglia, B., Bruzzone, M.G., Nardocci, N., Savoiardo, M., Chiapparini, L., 2009. Age-related Iron Deposition in the Basal Ganglia: Quantitative analysis in healthy subjects. Radiology 252, 165–172.

Ashby, F.G., 1983. A biased random walk model for two choice reaction times. J. Math. Psychol. 27, 277–297. https://doi.org/10.1016/0022-2496(83)90011-1

Avants, B., Epstein, C., Grossman, M., Gee, J., 2008. Symmetric diffeomorphic image registration with cross-correlation: Evaluating automated labeling of elderly and neurodegenerative brain. Med. Image Anal. 12, 26–41. https://doi.org/10.1016/j.media.2007.06.004

Ball, K., Sekuler, R., 1982. A specific and enduring improvement in visual motion discrimination. Science (80-.). 218, 697–698. https://doi.org/10.1126/science.7134968

Barr, D.J., Levy, R., Scheepers, C., Tily, H.J., 2013. Random effects structure for confirmatory hypothesis testing: Keep it maximal. J. Mem. Lang. 68, 255–278. https://doi.org/10.1016/j.jml.2012.11.001

Bates, D., Mächler, M., Bolker, B., Walker, S., 2015. Fitting Linear Mixed-Effects Models Using lme4. J. Stat. Softw. 67. https://doi.org/10.18637/jss.v067.i01

Behzadi, Y., Restom, K., Liau, J., Liu, T.T., 2007. A component based noise correction method (CompCor) for BOLD and perfusion based fMRI. Neuroimage 37, 90–101. https://doi.org/10.1016/j.neuroimage.2007.04.042

Binder, J.R., Liebenthal, E., Possing, E.T., Medler, D.A., Ward, B.D., 2004. Neural correlates of sensory and decision processes in auditory object identification. Nat. Neurosci. 7, 295–301. https://doi.org/10.1038/nn1198

Boehm, U., Annis, J., Frank, M.J., Hawkins, G.E., Heathcote, A., Kellen, D., Krypotos, A.M., Lerche, V., Logan, G.D., Palmeri, T.J., van Ravenzwaaij, D., Servant, M., Singmann, H., Starns, J.J., Voss, A., Wiecki, T. V., Matzke, D., Wagenmakers, E.J., 2018a. Estimating across-trial variability parameters of the Diffusion Decision Model: Expert advice and recommendations. J. Math. Psychol. 87, 46–75. https://doi.org/10.1016/j.jmp.2018.09.004

Boehm, U., Marsman, M., Matzke, D., Wagenmakers, E.-J., 2018b. On the importance of avoiding shortcuts in applying cognitive models to hierarchical data. Behav. Res. Methods 50, 1614–1631. https://doi.org/10.3758/s13428-018-1054-3

Bogacz, R., 2007. Optimal decision-making theories: linking neurobiology with behaviour. Trends Cogn. Sci. 11, 118–125. https://doi.org/10.1016/j.tics.2006.12.006

Bogacz, R., Gurney, K., 2007. The basal ganglia and cortex implement optimal decision making between alternative actions. Neural Comput. 19, 442–477. https://doi.org/10.1162/neco.2007.19.2.442

Boillat, Y., Van der Zwaag, W., 2019. Whole brain measurements of the positive BOLD response variability during a finger tapping task at 7 T show regional differences in its profiles. Magn. Reson. Med. 81, 2720–2727. https://doi.org/10.1002/mrm.27566

Botvinick, M.M., Braver, T.S., Barch, D.M., Carter, C.S., Cohen, J.D., 2001. Conflict monitoring and cognitive control. Psychol. Rev. 108, 624–652. https://doi.org/10.1037/0033-295X.108.3.624

Britten, K.H., Shadlen, M.N., Newsome, W.T., Movshon, J.A., 1993. Responses of neurons in macaque MT to stochastic motion signals. Vis. Neurosci. https://doi.org/10.1017/S0952523800010269

Britten, K.H., Shadlen, M.N., Newsome, W.T., Movshon, J.A., 1992. The analysis of visual motion: a comparison of neuronal and psychophysical performance. J. Neurosci. 12, 4745–4765. https://doi.org/10.1.1.123.9899

Brooks, S.P., Gelman, A., 1998. General Methods for Monitoring Convergence of Iterative Simulations. J. Comput. Graph. Stat. 7, 434–455. https://doi.org/10.1080/10618600.1998.10474787

Bullmore, E., Brammer, M.J., Williams, S.C.R., Rabe-Hesketh, S., Janot, N., David, A., Mellers, J., Howard, R., Sham, P., 1996. Statistical methods of estimation and inference for functional MR image analysis. Magn. Reson. Med. 35, 261–277. https://doi.org/10.1002/mrm.1910350219

Caan, M.W.A., Bazin, P.-L., Marques, J.P., Hollander, G., Dumoulin, S.O., Zwaag, W., 2019. MP2RAGEME: T1, T_2_*, and QSM mapping in one sequence at 7 tesla. Hum. Brain Mapp. 40, 1786–1798. https://doi.org/10.1002/hbm.24490

Christen, M., Bittlinger, M., Walter, H., Brugger, P., Müller, S., 2012. Dealing With Side Effects of Deep Brain Stimulation: Lessons Learned From Stimulating the STN. AJOB Neurosci. 3, 37–43. https://doi.org/10.1080/21507740.2011.635627

Coizet, V., Graham, J.H., Moss, J., Bolam, J.P., Savasta, M., McHaffie, J.G., Redgrave, P., Overton, P.G., 2009. Short-latency visual input to the subthalamic nucleus is provided by the midbrain superior colliculus. J. Neurosci. 29, 5701–5709. https://doi.org/10.1523/JNEUROSCI.0247-09.2009

Cousineau, D., 2005. Confidence intervals in within-subject designs: A simpler solution to Loftus and Masson's method. Tutor. Quant. Methods Psychol. 1, 42–45. https://doi.org/10.20982/tqmp.01.1.p042

Cox, R.W., Hyde, J.S., 1997. Software tools for analysis and visualization of fMRI data. NMR Biomed. 10, 171–178. https://doi.org/10.1002/(SICI)1099-1492(199706/08)10:4/5<171::AID-NBM453>3.0.CO;2-L

Cunnington, R., Bradshaw, J.L., Iansek, R., 1996. The role of the supplementary motor area in the control of voluntary movement. Hum. Mov. Sci. 15, 627–647. https://doi.org/10.1016/0167-9457(96)00018-8

Cunnington, R., Windischberger, C., Deecke, L., Moser, E., 2002. The preparation and execution of self-initiated and externally-triggered movement: A study of event-related fMRI. Neuroimage 15, 373–385. https://doi.org/10.1006/nimg.2001.0976

Cunnington, R., Windischberger, C., Moser, E., 2005. Premovement activity of the pre-supplementary motor area and the readiness for action: Studies of time-resolved event-related functional MRI. Hum. Mov. Sci. 24, 644–656. https://doi.org/10.1016/j.humov.2005.10.001

Dale, A.M., 1999. Optimal experimental design for event‐related fMRI. Hum. Brain Mapp. 8, 109–114. https://doi.org/10.1002/(SICI)1097-0193(1999)8:2/3<109::AID-HBM7>3.3.CO;2-N

Dale, A.M., Fischl, B., Sereno, M.I., 1999. Cortical Surface-Based Analysis. Neuroimage 9, 179–194. https://doi.org/10.1006/nimg.1998.0395

Dassonville, P., Zhu, X.H., Uǧurbil, K., Kim, S.-G., Ashe, J., 1997. Functional activation in motor cortex reflects the direction and the degree of handedness. Proc. Natl. Acad. Sci. U. S. A. 94, 14015–14018. https://doi.org/10.1073/pnas.94.25.14015

de Hollander, G., Forstmann, B.U., Brown, S.D., 2016. Different Ways of Linking Behavioral and Neural Data via Computational Cognitive Models. Biol. Psychiatry Cogn. Neurosci. Neuroimaging 1, 101–109. https://doi.org/10.1016/j.bpsc.2015.11.004

de Hollander, G., Keuken, M.C., Bazin, P.-L., Weiss, M., Neumann, J., Reimann, K., Wähnert, M., Turner, R., Forstmann, B.U., Schäfer, A., 2014. A gradual increase of iron toward the medial-inferior tip of the subthalamic nucleus. Hum. Brain Mapp. 35, 4440–4449. https://doi.org/10.1002/hbm.22485

de Hollander, G., Keuken, M.C., Forstmann, B.U., 2015. The subcortical cocktail problem; Mixed signals from the subthalamic nucleus and substantia nigra. PLoS One 10. https://doi.org/10.1371/journal.pone.0120572

de Hollander, G., Keuken, M.C., van der Zwaag, W., Forstmann, B.U., Trampel, R., 2017. Comparing functional MRI protocols for small, iron-rich basal ganglia nuclei such as the subthalamic nucleus at 7 T and 3 T. Hum. Brain Mapp. 38, 3226–3248. https://doi.org/10.1002/hbm.23586

de Hollander, G., Knapen, T., Snoek, L., 2019. Nideconv. http://doi.org/10.5281/zenodo.3240287

de Solages, C., Hill, B.C., Yu, H., Henderson, J.M., Bronte-Stewart, H., 2011. Maximal subthalamic beta hypersynchrony of the local field potential in Parkinson's disease is located in the central region of the nucleus. J. Neurol. Neurosurg. Psychiatry 82, 1387–1389. https://doi.org/10.1136/jnnp.2010.223107

Deistung, A., Schäfer, A., Schweser, F., Biedermann, U., Turner, R., Reichenbach, J.R., 2013. Toward in vivo histology: A comparison of quantitative susceptibility mapping (QSM) with magnitude-, phase-, and R2*-imaging at ultra-high magnetic field strength. Neuroimage 65, 299–314. https://doi.org/10.1016/j.neuroimage.2012.09.055

Devos, D., Szurhaj, W., Reyns, N., Labyt, E., Houdayer, E., Bourriez, J.L., Cassim, F., Krystkowiak, P., Blond, S., Destée, A., Derambure, P., Defebvre, L., 2006. Predominance of the contralateral movement-related activity in the subthalamo-cortical loop. Clin. Neurophysiol. 117, 2315–2327. https://doi.org/10.1016/j.clinph.2006.06.719

Diederich, A., Busemeyer, J.R., 2006. Modeling the effects of payoff on response bias in a perceptual discrimination task: Bound-change, drift-rate-change, or two-stage-processing hypothesis. Percept. Psychophys. 68, 194–207. https://doi.org/10.3758/BF03193669

Ditterich, J., 2006. Stochastic models of decisions about motion direction: Behavior and physiology. Neural Networks 19, 981–1012. https://doi.org/10.1016/j.neunet.2006.05.042

Donkin, C., Brown, S.D., Heathcote, A., 2009. The overconstraint of response time models: Rethinking the scaling problem. Psychon. Bull. Rev. 16, 1129–1135. https://doi.org/10.3758/PBR.16.6.1129

Duncan, J., Owen, A.M., 2000. Common regions of the human frontal lobe recruited by diverse cognitive demands. Trends Neurosci. 23, 475–483. https://doi.org/10.1016/S0166-2236(00)01633-7

Eckert, M.A., Menon, V., Walczak, A., Ahlstrom, J., Denslow, S., Horwitz, A., Dubno, J.R., 2009. At the heart of the ventral attention system: The right anterior insula. Hum. Brain Mapp. 30, 2530–2541. https://doi.org/10.1002/hbm.20688

Edwards, W., 1965. Optimal strategies for seeking information: Models for statistics, choice reaction times, and human information processing. J. Math. Psychol. 2, 312–329. https://doi.org/10.1016/0022-2496(65)90007-6

Emmi, A., Antonini, A., Macchi, V., Porzionato, A., De Caro, R., 2020. Anatomy and Connectivity of the Subthalamic Nucleus in Humans and Non-human Primates. Front. Neuroanat. 14. https://doi.org/10.3389/fnana.2020.00013

Esteban, O., Blair, R.W., Markiewicz, C.J., Berleant, S.L., Moodie, C., Ma, F., Isik, A.I., Erramuzpe, A., Kent, J.D., Goncalves, M., DuPre, E., Sitek, K.R., Gomez, D.E.P., Lurie, D.J., Ye, Z., Poldrack, R.A., Gorgolewski, K.J., 2018. fMRIPrep. Software. https://doi.org/10.5281/zenodo.852659

Esteban, O., Markiewicz, C.J., Blair, R.W., Moodie, C.A., Isik, A.I., Erramuzpe, A., Kent, J.D., Goncalves, M., DuPre, E., Snyder, M., Oya, H., Ghosh, S.S., Wright, J., Durnez, J., Poldrack, R.A., Gorgolewski, K.J., 2019. fMRIPrep: a robust preprocessing pipeline for functional MRI. Nat. Methods 16, 111–116. https://doi.org/10.1038/s41592-018-0235-4

Evans, N.J., Hawkins, G.E., 2019. When humans behave like monkeys: Feedback delays and extensive practice increase the efficiency of speeded decisions. Cognition 184, 11–18. https://doi.org/10.1016/j.cognition.2018.11.014

Ewert, S., Plettig, P., Li, N., Chakravarty, M.M., Collins, D.L., Herrington, T.M., Kühn, A.A., Horn, A., 2018. Toward defining deep brain stimulation targets in MNI space: A subcortical atlas based on multimodal MRI, histology and structural connectivity. Neuroimage 170, 271–282. https://doi.org/10.1016/j.neuroimage.2017.05.015

Fasano, A., Lozano, A.M., 2015. Deep brain stimulation for movement disorders. Curr. Opin. Neurol. 28, 423–436. https://doi.org/10.1097/WCO.0000000000000226

Fonov, V., Evans, A., McKinstry, R., Almli, C., Collins, D., 2009. Unbiased nonlinear average age-appropriate brain templates from birth to adulthood. Neuroimage 47, S102. https://doi.org/10.1016/S1053-8119(09)70884-5

Forstmann, B.U., Anwander, A., Schafer, A., Neumann, J., Brown, S., Wagenmakers, E.-J., Bogacz, R., Turner, R., 2010a. Cortico-striatal connections predict control over speed and accuracy in perceptual decision making. Proc. Natl. Acad. Sci. 107, 15916–15920. https://doi.org/10.1073/pnas.1004932107

Forstmann, B.U., Brown, S.D., Dutilh, G., Neumann, J., Wagenmakers, E.-J., 2010b. The neural substrate of prior information in perceptual decision making: a model-based analysis. Front. Hum. Neurosci. 4, 1–12. https://doi.org/10.3389/fnhum.2010.00040

Forstmann, B.U., de Hollander, G., Van Maanen, L., Alkemade, A., Keuken, M.C., 2017. Towards a mechanistic understanding of the human subcortex. Nat. Rev. Neurosci. 18, 57–65.

Forstmann, B.U., Dutilh, G., Brown, S.D., Neumann, J., von Cramon, D.Y., Ridderinkhof, K.R., Wagenmakers, E.-J., 2008. Striatum and pre-SMA facilitate decision-making under time pressure. Proc. Natl. Acad. Sci. U. S. A. 105, 17538–17542. https://doi.org/10.1073/pnas.0805903105

Forstmann, B.U., Ratcliff, R., Wagenmakers, E.-J., 2016. Sequential Sampling Models in Cognitive Neuroscience: Advantages, Applications, and Extensions. Annu. Rev. Psychol. 67, 641–666. https://doi.org/10.1146/annurev-psych-122414-033645

Forstmann, B.U., Wagenmakers, E., Eichele, T., Brown, S.D., Serences, J.T., 2011. Reciprocal relations between cognitive neuroscience and formal cognitive models: opposites attract? Trends Cogn. Sci. 15, 272–279. https://doi.org/10.1016/j.tics.2011.04.002

Frank, M.J., 2006. Hold your horses: A dynamic computational role for the subthalamic nucleus in decision making. Neural Networks 19, 1120–1136. https://doi.org/10.1016/j.neunet.2006.03.006

Frank, M.J., Gagne, C., Nyhus, E., Masters, S., Wiecki, T. V., Cavanagh, J.F., Badre, D., 2015. fMRI and EEG Predictors of Dynamic Decision Parameters during Human Reinforcement Learning. J. Neurosci. 35, 485–494. https://doi.org/10.1523/JNEUROSCI.2036-14.2015

Gelman, A., Hill, J., 2007. Data Analysis Using Regression and Multilevel/Hierarchical Models. Cambridge University Press, Cambridge.

Gelman, A., Rubin, D.B., 1992. Inference from Iterative Simulation Using Multiple Sequences. Stat. Sci. 7, 457–472. https://doi.org/10.1214/ss/1177011136

Gitelman, D.R., Penny, W.D., Ashburner, J., Friston, K.J., 2003. Modeling regional and psychophysiologic interactions in fMRI: The importance of hemodynamic deconvolution. Neuroimage 19, 200–207. https://doi.org/10.1016/S1053-8119(03)00058-2

Glasser, M.F., Sotiropoulos, S.N., Wilson, J.A., Coalson, T.S., Fischl, B., Andersson, J.L.R., Xu, J., Jbabdi, S., Webster, M., Polimeni, J.R., Van Essen, D.C., Jenkinson, M., 2013. The minimal preprocessing pipelines for the Human Connectome Project. Neuroimage 80, 105–124. https://doi.org/10.1016/j.neuroimage.2013.04.127

Glover, G.H., 1999. Deconvolution of Impulse Response in Event-Related BOLD fMRI. Neuroimage 9, 416–429.

Gold, J.I., Shadlen, M.N., 2001. Neural computations that underlie decisions about sensory stimuli. Trends Cogn. Sci. 5, 10–16. https://doi.org/10.1016/S1364-6613(00)01567-9

Gonzalez-Castillo, J., Saad, Z.S., Handwerker, D.A., Inati, S.J., Brenowitz, N., Bandettini, P.A., 2012. Whole-brain, time-locked activation with simple tasks revealed using massive averaging and model-free analysis. Proc. Natl. Acad. Sci. U. S. A. 109, 5487–5492. https://doi.org/10.1073/pnas.1121049109

Gorgolewski, K.J., Burns, C.D., Madison, C., Clark, D., Halchenko, Y.O., Waskom, M.L., Ghosh, S.S., 2011. Nipype: A Flexible, Lightweight and Extensible Neuroimaging Data Processing Framework in Python. Front. Neuroinform. 5. https://doi.org/10.3389/fninf.2011.00013

Gorgolewski, K.J., Esteban, O., Markiewicz, C.J., Ziegler, E., Ellis, D.G., Notter, M.P., Jarecka, D., Johnson, H., Burns, C.D., Manhães-Savio, A., Hamalainen, C., Yvernault, B., Salo, T., Jordan, K., Goncalves, M., Waskom, M., Clark, Daniel, Wong, J., Loney, F., Modat, M., Dewey, B.E., Madison, C., Visconti di Oleggio Castello, M., Clark, M.G., Dayan, M., Clark, Dav, Keshavan, A., Pinsard, B., Gramfort, A., Berleant, S.L., Nielson, D.M., Bougacha, S., Varoquaux, G., Cipollini, B., Markello, R., Rokem, A., Moloney, B., Halchenko, Y.O., Wassermann, D., Hanke, M., Horea, C., Kaczmarzyk, J., Hollander, G. de, DuPre, E., Gillman, A., Mordom, D., Buchanan, C., Tungaraza, R., Pauli, W.M., Iqbal, S., Sikka, S., Mancini, M., Schwartz, Y., Malone, I.B., Dubois, M., Frohlich, C., Welch, D., Forbes, J., Kent, J., Watanabe, A., Cumba, C., Huntenburg, J.M., Kastman, E., Nichols, B.N., Eshaghi, A., Ginsburg, D., Schaefer, A., Acland, B., Giavasis, S., Kleesiek, J., Erickson, D., Küttner, R., Haselgrove, C., Correa, C., Ghayoor, A., Liem, F., Millman, J., Haehn, D., Lai, J., Zhou, D., Blair, R.W., Glatard, T., Renfro, M., Liu, S., Kahn, A.E., Pérez-García, F., Triplett, W., Lampe, L., Stadler, J., Kong, X.-Z., Hallquist, M., Chetverikov, A., Salvatore, J., Park, A., Poldrack, R., Craddock, R.C., Inati, S., Hinds, O., Cooper, G., Perkins, L.N., Marina, A., Mattfeld, A., Noel, M., Snoek, L., Matsubara, K., Cheung, B., Rothmei, S., Urchs, S., Durnez, J., Mertz, F., Geisler, D., Floren, A., Gerhard, S., Sharp, P., Molina-Romero, M., Weinstein, A., Broderick, W., Saase, V., Andberg, S.K., Harms, R., Schlamp, K., Arias, J., Papadopoulos Orfanos, D., Tarbert, C., Tambini, A., De La Vega, A., Nickson, T., Brett, M., Falkiewicz, M., Podranski, K., Linkersdörfer, J., Flandin, G., Ort, E., Shachnev, D., McNamee, D., Davison, A., Varada, J., Schwabacher, I., Pellman, J., Perez-Guevara, M., Khanuja, R., Pannetier, N., McDermottroe, C., Ghosh, S.S., 2018. Nipype. https://doi.org/10.5281/zenodo.596855

Green, N., Bogacz, R., Huebl, J., Beyer, A.K., Kühn, A.A., Heekeren, H.R., 2013. Reduction of influence of task difficulty on perceptual decision making by stn deep brain stimulation. Curr. Biol. 23, 1681–1684. https://doi.org/10.1016/j.cub.2013.07.001

Greenhouse, I., Gould, S., Houser, M., Aron, A.R., 2013. Stimulation of contacts in ventral but not dorsal subthalamic nucleus normalizes response switching in Parkinson's disease. Neuropsychologia 51, 1302–1309. https://doi.org/10.1016/j.neuropsychologia.2013.03.008

Greenhouse, I., Gould, S., Houser, M., Hicks, G., Gross, J., Aron, A.R., 2011. Stimulation at dorsal and ventral electrode contacts targeted at the subthalamic nucleus has different effects on motor and emotion functions in Parkinson's disease. Neuropsychologia 49, 528–534. https://doi.org/10.1016/j.neuropsychologia.2010.12.030

Greve, D.N., Fischl, B., 2009. Accurate and robust brain image alignment using boundary-based registration. Neuroimage 48, 63–72. https://doi.org/10.1016/j.neuroimage.2009.06.060

Groiss, S.J., Wojtecki, L., Südmeyer, M., Schnitzler, A., 2009. Review: Deep brain stimulation in Parkinson's disease. Ther. Adv. Neurol. Disord. 2, 379–391. https://doi.org/10.1177/1756285609339382

Handwerker, D.A., Ollinger, J.M., D’Esposito, M., 2004. Variation of BOLD hemodynamic responses across subjects and brain regions and their effects on statistical analyses. Neuroimage 21, 1639–1651. https://doi.org/10.1016/j.neuroimage.2003.11.029

Hawkins, G.E., Forstmann, B.U., Wagenmakers, E.-J., Ratcliff, R., Brown, S.D., 2015. Revisiting the Evidence for Collapsing Boundaries and Urgency Signals in Perceptual Decision-Making. J. Neurosci. 35, 2476–2484. https://doi.org/10.1523/JNEUROSCI.2410-14.2015

Haynes, W.I.A., Haber, S.N., 2013. The Organization of Prefrontal-Subthalamic Inputs in Primates Provides an Anatomical Substrate for Both Functional Specificity and Integration: Implications for Basal Ganglia Models and Deep Brain Stimulation. J. Neurosci. 33, 4804–4814. https://doi.org/10.1523/JNEUROSCI.4674-12.2013

Heathcote, A., Lin, Y.S., Reynolds, A., Strickland, L., Gretton, M., Matzke, D., 2019. Dynamic models of choice. Behav. Res. Methods 51, 961–985. https://doi.org/10.3758/s13428-018-1067-y

Heekeren, H.R., Marrett, S., Bandettini, P. a., Ungerleider, L.G., 2004. A general mechanism for perceptual decision-making in the human brain. Nature 431, 859–862. https://doi.org/10.1038/nature02966

Herz, D.M., Zavala, B.A., Bogacz, R., Brown, P., 2016. Neural Correlates of Decision Thresholds in the Human Subthalamic Nucleus. Curr. Biol. 26, 916–920. https://doi.org/10.1016/j.cub.2016.01.051

Ho, T.C., Brown, S.D., Serences, J.T., 2009. Domain General Mechanisms of Perceptual Decision Making in Human Cortex. J. Neurosci. 29, 8675–8687. https://doi.org/10.1523/JNEUROSCI.5984-08.2009

Hoeting, J.J.A., Madigan, D., Raftery, A.E.A., Volinsky, C.T., 1999. Bayesian model averaging: A tutorial. Stat. Sci. 14, 382–401. https://doi.org/10.2307/2676803

Horn, A., Neumann, W.-J., Degen, K., Schneider, G.-H., Kühn, A.A., 2017. Toward an electrophysiological “sweet spot” for deep brain stimulation in the subthalamic nucleus. Hum. Brain Mapp. https://doi.org/10.1002/hbm.23594

Huber, L., Goense, J., Kennerley, A.J., Trampel, R., Guidi, M., Reimer, E., Ivanov, D., Neef, N., Gauthier, C.J., Turner, R., Möller, H.E., 2015. Cortical lamina-dependent blood volume changes in human brain at 7T. Neuroimage 107, 23–33. https://doi.org/10.1016/j.neuroimage.2014.11.046

Huber, L., Ivanov, D., Handwerker, D.A., Marrett, S., Guidi, M., Uludaǧ, K., Bandettini, P.A., Poser, B.A., 2018. Techniques for blood volume fMRI with VASO: From low-resolution mapping towards sub-millimeter layer-dependent applications. Neuroimage 164, 131–143. https://doi.org/10.1016/j.neuroimage.2016.11.039

Huber, L., Ivanov, D., Krieger, S.N., Streicher, M.N., Mildner, T., Poser, B.A., Möller, H.E., Turner, R., 2014. Slab-selective, BOLD-corrected VASO at 7 tesla provides measures of cerebral blood volume reactivity with high signal-to-noise ratio. Magn. Reson. Med. 72, 137–148. https://doi.org/10.1002/mrm.24916

Huber, L., Uludaǧ, K., Möller, H.E., 2019. Non-BOLD contrast for laminar fMRI in humans: CBF, CBV, and CMRO2. Neuroimage 197, 742–760. https://doi.org/10.1016/j.neuroimage.2017.07.041

Huntenburg, J.M., 2014. Evaluating Nonlinear Coregistration of BOLD EPI and T1w Images. Freie Universität, Berlin.

Jeffreys, H., 1961. The theory of probability, 3rd ed. Oxford University Press, Oxford.

Jenkinson, M., Bannister, P., Brady, M., Smith, S., 2002. Improved Optimization for the Robust and Accurate Linear Registration and Motion Correction of Brain Images. Neuroimage 17, 825–841. https://doi.org/10.1006/nimg.2002.1132

Joel, D., Weiner, I., 1997. The connections of the primate subthalamic nucleus: indirect pathways and the open-interconnected scheme of basal ganglia-thalamocortical circuitry. Brain Res. Rev. 23, 62–78. https://doi.org/10.1016/S0165-0173(96)00018-5

Kaiser, J., Lennert, T., Lutzenberger, W., 2007. Dynamics of oscillatory activity during auditory decision making. Cereb. Cortex 17, 2258–2267. https://doi.org/10.1093/cercor/bhl134

Karachi, C., Grabli, D., Baup, N., Mounayar, S., Tandé, D., François, C., Hirsch, E.C., 2009. Dysfunction of the subthalamic nucleus induces behavioral and movement disorders in monkeys. Mov. Disord. 24, 1183–1192. https://doi.org/10.1002/mds.22547

Kass, B., Raftery, A., 1995. Bayes Factors. J. Am. Stat. Assoc. 90(430), pp.773–795.

Katsimpokis, D., Hawkins, G.E., Van Maanen, L., 2020. Not all Speed-Accuracy Trade-Off Manipulations Have the Same Psychological Effect. Comput. Brain Behav. https://doi.org/10.1007/s42113-020-00074-y

Keuken, M.C., Bazin, P., Backhouse, K., Beekhuizen, S., Himmer, L., Kandola, A., Lafeber, J.J., Prochazkova, L., Trutti, A., Schäfer, A., Turner, R., Forstmann, B.U., 2017. Effects of aging on T1, T2∗, and QSM MRI values in the subcortex. Brain Struct. Funct. 222, 2487–2505. https://doi.org/10.1007/s00429-016-1352-4

Keuken, M.C., Bazin, P., Crown, L., Hootsmans, J., Laufer, A., Müller-Axt, C., Sier, R., van der Putten, E.J., Schäfer, A., Turner, R., Forstmann, B.U., 2014a. Quantifying inter-individual anatomical variability in the subcortex using 7T structural MRI. Neuroimage 94, 40–46. https://doi.org/10.1016/j.neuroimage.2014.03.032

Keuken, M.C., Isaacs, B.R., Trampel, R., van der Zwaag, W., Forstmann, B.U., 2018a. Visualizing the Human Subcortex Using Ultra-high Field Magnetic Resonance Imaging. Brain Topogr. 31, 513–545. https://doi.org/10.1007/s10548-018-0638-7

Keuken, M.C., Mü, Ller-Axt, C., Langner, R., Eickhoff, S.B., Forstmann, B.U., Neumann, J., 2014b. Brain networks of perceptual decision-making: an fMRI ALE meta-analysis. Front. Hum. Neurosci. 8. https://doi.org/10.3389/fnhum.2014.00445

Keuken, M.C., Uylings, H.B.M., Geyer, S., Schäfer, A., Turner, R., Forstmann, B.U., 2012. Are there three subdivisions in the primate subthalamic nucleus? Front. Neuroanat. 6. https://doi.org/10.3389/fnana.2012.00014

Keuken, M.C., Van Maanen, L., Bogacz, R., Schäfer, A., Neumann, J., Turner, R., Forstmann, B.U., 2015. The subthalamic nucleus during decision-making with multiple alternatives. Hum. Brain Mapp. 36, 4041–4052. https://doi.org/10.1002/hbm.22896

Keuken, M.C., van Maanen, L., Boswijk, M., Forstmann, B.U., Steyvers, M., 2018b. Large scale structure-function mappings of the human subcortex. Sci. Rep. 8, 15854. https://doi.org/10.1038/s41598-018-33796-y

Kim, J.N., Shadlen, M.N., 1999. Neural correlates of a decision in the dorsolateral prefrontal cortex of the macaque. Nat. Neurosci. 2, 176–185. https://doi.org/10.1038/5739

Kim, S.-G., Ashe, J., Georgopoulos, A.P., Merkle, H., Ellermann, J.M., Menon, R.S., Ogawa, S., Ugurbil, K., 1993a. Functional imaging of human motor cortex at high magnetic field. J. Neurophysiol. 69, 297–302. https://doi.org/10.1152/jn.1993.69.1.297

Kim, S.-G., Ashe, J., Hendrich, K., Ellermann, J., Merkle, H., Ugurbil, K., Georgopoulos, A.P., 1993b. Functional magnetic resonance imaging of motor cortex: hemispheric asymmetry and handedness. Science (80-.). 261, 615–617. https://doi.org/10.1126/science.8342027

Klein, A., Ghosh, S.S., Bao, F.S., Giard, J., Häme, Y., Stavsky, E., Lee, N., Rossa, B., Reuter, M., Chaibub Neto, E., Keshavan, A., 2017. Mindboggling morphometry of human brains. PLOS Comput. Biol. 13, e1005350. https://doi.org/10.1371/journal.pcbi.1005350

Klein, T.A., Ullsperger, M., Danielmeier, C., 2013. Error awareness and the insula: Links to neurological and psychiatric diseases. Front. Hum. Neurosci. 7, 1–14. https://doi.org/10.3389/fnhum.2013.00014

Kühn, A.A., Trottenberg, T., Kivi, A., Kupsch, A., Schneider, G.-H., Brown, P., 2005. The relationship between local field potential and neuronal discharge in the subthalamic nucleus of patients with Parkinson's disease. Exp. Neurol. 194, 212–220. https://doi.org/10.1016/j.expneurol.2005.02.010

Kundu, P., Inati, S.J., Evans, J.W., Luh, W.M., Bandettini, P.A., 2012. Differentiating BOLD and non-BOLD signals in fMRI time series using multi-echo EPI. Neuroimage 60, 1759–1770. https://doi.org/10.1016/j.neuroimage.2011.12.028

Kuznetsova, A., Brockhoff, P.B., Christensen, R.H.B., 2017. lmerTest Package: Tests in Linear Mixed Effects Models. J. Stat. Softw. 82. https://doi.org/10.18637/jss.v082.i13

Lambert, C., Zrinzo, L., Nagy, Z., Lutti, A., Hariz, M., Foltynie, T., Draganski, B., Ashburner, J., Frackowiak, R., 2015. Do we need to revise the tripartite subdivision hypothesis of the human subthalamic nucleus (STN)? Response to Alkemade and Forstmann. Neuroimage 110, 1–2. https://doi.org/10.1016/j.neuroimage.2015.01.038

Lambert, C., Zrinzo, L., Nagy, Z., Lutti, A., Hariz, M., Foltynie, T., Draganski, B., Ashburner, J., Frackowiak, R., 2012. Confirmation of functional zones within the human subthalamic nucleus: Patterns of connectivity and sub-parcellation using diffusion weighted imaging. Neuroimage 60, 83–94. https://doi.org/10.1016/j.neuroimage.2011.11.082

Lanczos, C., 1964. Evaluation of Noisy Data. J. Soc. Ind. Appl. Math. Ser. B Numer. Anal. 1, 76–85. https://doi.org/10.1137/0701007

Lebreton, M., Bavard, S., Daunizeau, J., Palminteri, S., 2019. Assessing inter-individual differences with task-related functional neuroimaging. Nat. Hum. Behav. 3, 897–905. https://doi.org/10.1038/s41562-019-0681-8

Leech, R., Sharp, D.J., 2014. The role of the posterior cingulate cortex in cognition and disease. Brain 137, 12–32. https://doi.org/10.1093/brain/awt162

Limousin, P., Pollak, P., Benazzouz, A., Hoffmann, D., Le Bas, J.-F., Perret, J.E., Benabid, A.-L., Broussolle, E., 1995. Effect on parkinsonian signs and symptoms of bilateral subthalamic nucleus stimulation. Lancet 345, 91–95. https://doi.org/10.1016/S0140-6736(95)90062-4

Link, S.W., Heath, R.A., 1975. A sequential theory of psychological discrimination. Psychometrika 40, 77–105. https://doi.org/10.1007/BF02291481

Lotze, M., Montoya, P., Erb, M., Hülsmann, E., Flor, H., Klose, U., Birbaumer, N., Grodd, W., 1999. Activation of cortical and cerebellar motor areas during executed and imagined hand movements: An fMRI study. J. Cogn. Neurosci. 11, 491–501. https://doi.org/10.1162/089892999563553

Lozano, A.M., Lipsman, N., 2013. Probing and Regulating Dysfunctional Circuits Using Deep Brain Stimulation. Neuron 77, 406–424. https://doi.org/10.1016/j.neuron.2013.01.020

Ly, A., Verhagen, J., Wagenmakers, E.J., 2016. Harold Jeffreys's default Bayes factor hypothesis tests: Explanation, extension, and application in psychology. J. Math. Psychol. 72, 19–32. https://doi.org/10.1016/j.jmp.2015.06.004

Mallet, L., Schupbach, M., N’Diaye, K., Remy, P., Bardinet, E., Czernecki, V., Welter, M.-L., Pelissolo, A., Ruberg, M., Agid, Y., Yelnik, J., 2007. Stimulation of subterritories of the subthalamic nucleus reveals its role in the integration of the emotional and motor aspects of behavior. Proc. Natl. Acad. Sci. 104, 10661–10666. https://doi.org/10.1073/pnas.0610849104

Markuerkiaga, I., Barth, M., Norris, D.G., 2016. A cortical vascular model for examining the specificity of the laminar BOLD signal. Neuroimage 132, 491–498. https://doi.org/10.1016/j.neuroimage.2016.02.073

Marques, J.P., Kober, T., Krueger, G., van der Zwaag, W., Van de Moortele, P.F., Gruetter, R., 2010. MP2RAGE, a self bias-field corrected sequence for improved segmentation and T1-mapping at high field. Neuroimage 49, 1271–1281. https://doi.org/10.1016/j.neuroimage.2009.10.002

McClure, S.M., Berns, G.S., Montague, P.R., 2003. Temporal prediction errors in a passive learning task activate human striatum. Neuron 38, 339–346. https://doi.org/10.1016/S0896-6273(03)00154-5

Miezin, F.M., Maccotta, L., Ollinger, J.M., Petersen, S.E., Buckner, R.L., 2000. Characterizing the hemodynamic response: Effects of presentation rate, sampling procedure, and the possibility of ordering brain activity based on relative timing. Neuroimage 11, 735–759. https://doi.org/10.1006/nimg.2000.0568

Miletić, S., Bazin, P.-L., Weiskopf, N., van der Zwaag, W., Forstmann, B.U., Trampel, R., 2020. fMRI protocol optimization for simultaneously studying small subcortical and cortical areas at 7 T. Neuroimage 219. https://doi.org/10.1016/j.neuroimage.2020.116992

Miletić, S., Van Maanen, L., 2019. Caution in decision-making under time pressure is mediated by timing ability. Cogn. Psychol. 110, 16–29. https://doi.org/10.1016/j.cogpsych.2019.01.002

Mink, J.W., 1996. The Basal Ganglia: Focused selection and inhibition of competing motor programs. Prog. Neurobiol. 50, 381–425. https://doi.org/10.1016/S0301-0082(96)00042-1

Morey, R.D., Rouder, J.N., Jamil, T., Urbanek, S., Forner, K., Ly, A., 2018. BayesFactor.

Mulder, M.J., Keuken, M.C., Bazin, P., Alkemade, A., Forstmann, B.U., 2019. Size and shape matter: The impact of voxel geometry on the identification of small nuclei. PLoS One 14, e0215382. https://doi.org/10.1371/journal.pone.0215382

Mulder, M.J., van Maanen, L., Forstmann, B.U., 2014. Perceptual decision neurosciences – A model-based review. Neuroscience 277, 872–884. https://doi.org/10.1016/j.neuroscience.2014.07.031

Mulder, M.J., Wagenmakers, E.-J., Ratcliff, R., Boekel, W., Forstmann, B.U., 2012. Bias in the Brain: A Diffusion Model Analysis of Prior Probability and Potential Payoff. J. Neurosci. 32, 2335–2343. https://doi.org/10.1523/JNEUROSCI.4156-11.2012

Nieuwenhuis, S., Forstmann, B.U., Wagenmakers, E.-J., 2011. Erroneous analyses of interactions in neuroscience: a problem of significance. Nat. Neurosci. 14, 1105–7. https://doi.org/10.1038/nn.2886

O’Connell, R.G., Dockree, P.M., Kelly, S.P., 2012. A supramodal accumulation-to-bound signal that determines perceptual decisions in humans. Nat. Neurosci. 15, 1729–1735. https://doi.org/10.1038/nn.3248

Oldfield, R.C., 1971. The assessment and analysis of handedness: The Edinburgh inventory. Neuropsychologia 9, 97–113. https://doi.org/10.1016/0028-3932(71)90067-4

Palmer, J., Huk, A.C., Shadlen, M.N., 2005. The effect of stimulus strength on the speed and accuracy of a perceptual decision. J. Vis. 5, 1. https://doi.org/10.1167/5.5.1

Parent, A., Hazrati, L.-N., 1995. Functional anatomy of the basal ganglia. II. The place of subthalamic nucleus and external pallidium in basal ganglia circuitry. Brain Res. Rev. 20, 128–154. https://doi.org/10.1016/0165-0173(94)00008-D

Paus, T., 2001. Primate anterior cingulate cortex: Where motor control, drive and cognition interface. Nat. Rev. Neurosci. 2, 417–424. https://doi.org/10.1038/35077500

Paus, T., Koski, L., Caramanos, Z., Westbury, C., 1998. Regional differences in the effects of task difficulty and motor output on blood flow response in the human anterior cingulate cortex: A review of 107 PET activation studies. Neuroreport. https://doi.org/10.1097/00001756-199806220-00001

Pedersen, J.R., Johannsen, P., Bak, C.K., Kofoed, B., Saermark, K., Gjedde, A., 1998. Origin of human motor Readiness Field linked to left middle frontal gyrus by MEG and PET. Neuroimage 8, 214–220. https://doi.org/10.1006/nimg.1998.0362

Pilly, P.K., Seitz, A.R., 2009. What a difference a parameter makes: A psychophysical comparison of random dot motion algorithms. Vision Res. 49, 1599–1612. https://doi.org/10.1016/j.visres.2009.03.019

Poldrack, R.A., Mumford, J.A., Nichols, T.E., 2011. Statistical modeling: Single subject analysis, in: Handbook of Functional MRI. Cambridge University Press, Cambridge, pp. 70–99.

Posse, S., Wiese, S., Gembris, D., Mathiak, K., Kessler, C., Grosse-Ruyken, M.L., Elghahwagi, B., Richards, T., Dager, S.R., Kiselev, V.G., 1999. Enhancement of BOLD-Contrast Sensitivity by Single-Shot Multi-Echo Functional MR Imaging. Magn. Reson. Med. 42, 87–97.

Power, J.D., Mitra, A., Laumann, T.O., Snyder, A.Z., Schlaggar, B.L., Petersen, S.E., 2014. Methods to detect, characterize, and remove motion artifact in resting state fMRI. Neuroimage 84, 320–341. https://doi.org/10.1016/j.neuroimage.2013.08.048

R Core Team, 2017. R: A language and environment for statistical computing.

Ratcliff, R., 2006. Modeling response signal and response time data. Cogn. Psychol. 53, 195–237. https://doi.org/10.1016/j.cogpsych.2005.10.002

Ratcliff, R., 1985. Theoretical Interpretations of the Speed and Accuracy of Positive and Negative Responses. Psychol. Rev. 92, 212–225. https://doi.org/10.1037/0033-295X.92.2.212

Ratcliff, R., 1981. A theory of order relations in perceptual matching. Psychol. Rev. 88, 552–572. https://doi.org/10.1037/0033-295X.88.6.552

Ratcliff, R., 1978. A theory of memory retrieval. Psychol. Rev. 85, 59–108.

Ratcliff, R., Childers, R., 2015. Individual differences and fitting methods for the two-choice diffusion model of decision making. Decision 2, 237–279. https://doi.org/10.1037/dec0000030

Ratcliff, R., McKoon, G., 2008. The diffusion decision model: theory and data for two-choice decision tasks. Neural Comput. 20, 873–922. https://doi.org/10.1162/neco.2008.12-06-420

Ratcliff, R., Rouder, J.N., 1998. Modeling Response Times for Two-Choice Decisions. Psychol. Sci. 9, 347–356.

Ratcliff, R., Smith, P.L., Brown, S.D., McKoon, G., 2016. Diffusion Decision Model: Current Issues and History. Trends Cogn. Sci. 20, 260–281. https://doi.org/10.1016/j.tics.2016.01.007

Ratcliff, R., Tuerlinckx, F., 2002. Estimating parameters of the diffusion model: Approaches to dealing with contaminant reaction times and parameter variability. Psychon. Bull. Rev. 9, 438–481.

Reynolds, J.H., Heeger, D.J., 2009. The Normalization Model of Attention. Neuron 61, 168–185. https://doi.org/10.1016/j.neuron.2009.01.002

Satterthwaite, F.E., 1941. Synthesis of variance. Psychometrika 6, 309–316. https://doi.org/10.1007/BF02288586

Satterthwaite, T.D., Elliott, M.A., Gerraty, R.T., Ruparel, K., Loughead, J., Calkins, M.E., Eickhoff, S.B., Hakonarson, H., Gur, R.C., Gur, R.E., Wolf, D.H., 2013. An improved framework for confound regression and filtering for control of motion artifact in the preprocessing of resting-state functional connectivity data. Neuroimage 64, 240–256. https://doi.org/10.1016/j.neuroimage.2012.08.052

Seifried, C., Weise, L., Hartmann, R., Gasser, T., Baudrexel, S., Szelényi, A., van de Loo, S., Steinmetz, H., Seifert, V., Roeper, J., Hilker, R., 2012. Intraoperative microelectrode recording for the delineation of subthalamic nucleus topography in Parkinson's disease. Brain Stimul. 5, 378–387. https://doi.org/10.1016/j.brs.2011.06.002

Shadlen, M.N., Newsome, W.T., 2001. Neural basis of a perceptual decision in the parietal cortex (area LIP) of the rhesus monkey. J. Neurophysiol. 86, 1916–36.

Shmuel, A., Yacoub, E., Chaimow, D., Logothetis, N.K., Ugurbil, K., 2007. Spatio-temporal point-spread function of fMRI signal in human gray matter at 7 Tesla. Neuroimage 35, 539–552. https://doi.org/10.1016/j.neuroimage.2006.12.030

Smith, S.M., Brady, J.M., 1997. SUSAN—A New Approach to Low Level Image Processing. Int. J. Comput. Vis. 23, 45–78. https://doi.org/10.1023/A:1007963824710

Soh, C., Wessel, J.R., 2021. Unexpected Sounds Nonselectively Inhibit Active Visual Stimulus Representations. Cereb. Cortex 31, 1632–1646. https://doi.org/10.1093/cercor/bhaa315

Spunt, R.P., Lieberman, M.D., Cohen, J.R., Eisenberger, N.I., 2012. The phenomenology of error processing: The dorsal ACC response to stop-signal errors tracks reports of negative affect. J. Cogn. Neurosci. 24, 1753–1765. https://doi.org/10.1162/jocn_a_00242

Stüber, C., Morawski, M., Schäfer, A., Labadie, C., Wähnert, M., Leuze, C., Streicher, M., Barapatre, N., Reimann, K., Geyer, S., Spemann, D., Turner, R., 2014. Myelin and iron concentration in the human brain: A quantitative study of MRI contrast. Neuroimage 93, 95–106. https://doi.org/10.1016/j.neuroimage.2014.02.026

Summerfield, C., Koechlin, E., 2010. Economic Value Biases Uncertain Perceptual Choices in the Parietal and Prefrontal Cortices. Front. Hum. Neurosci. 4, 1–12. https://doi.org/10.3389/fnhum.2010.00208

Sylvester, R., Haynes, J.-D., Rees, G., 2005. Saccades Differentially Modulate Human LGN and V1 Responses in the Presence and Absence of Visual Stimulation. Curr. Biol. 15, 37–41. https://doi.org/10.1016/j.cub.2004.12.061

Temel, Y., Blokland, A., Steinbusch, H.W.M., Visser-Vandewalle, V., 2005a. The functional role of the subthalamic nucleus in cognitive and limbic circuits. Prog. Neurobiol. 76, 393–413. https://doi.org/10.1016/j.pneurobio.2005.09.005

Temel, Y., Kessels, A., Tan, S., Topdag, A., Boon, P., Visser-Vandewalle, V., 2006. Behavioural changes after bilateral subthalamic stimulation in advanced Parkinson disease: A systematic review. Parkinsonism Relat. Disord. 12, 265–272. https://doi.org/10.1016/j.parkreldis.2006.01.004

Temel, Y., Visser-Vandewalle, V., Aendekerk, B., Rutten, B., Tan, S., Scholtissen, B., Schmitz, C., Blokland, A., Steinbusch, H.W.M., 2005b. Acute and separate modulation of motor and cognitive performance in parkinsonian rats by bilateral stimulation of the subthalamic nucleus. Exp. Neurol. 193, 43–52. https://doi.org/10.1016/j.expneurol.2004.12.025

Ter Braak, C.J.F., 2006. A Markov Chain Monte Carlo version of the genetic algorithm Differential Evolution: easy Bayesian computing for real parameter spaces. Stat. Comput. 16, 239–249. https://doi.org/10.1007/s11222-006-8769-1

Thielscher, A., Pessoa, L., 2007. Neural correlates of perceptual choice and decision making during fear-disgust discrimination. J. Neurosci. 27, 2908–2917. https://doi.org/10.1523/JNEUROSCI.3024-06.2007

Treiber, J.M., White, N.S., Steed, T.C., Bartsch, H., Holland, D., Farid, N., McDonald, C.R., Carter, B.S., Dale, A.M., Chen, C.C., 2016. Characterization and Correction of Geometric Distortions in 814 Diffusion Weighted Images. PLoS One 11, e0152472. https://doi.org/10.1371/journal.pone.0152472

Trottenberg, T., Kupsch, A., Schneider, G., Brown, P., Kühn, A., 2007. Frequency-dependent distribution of local field potential activity within the subthalamic nucleus in Parkinson's disease. Exp. Neurol. 205, 287–291. https://doi.org/10.1016/j.expneurol.2007.01.028

Turner, B.M., Forstmann, B.U., Love, B.C., Palmeri, T.J., Van Maanen, L., 2017. Approaches to analysis in model-based cognitive neuroscience. J. Math. Psychol. 76, 65–79. https://doi.org/10.1016/j.jmp.2016.01.001

Turner, B.M., Palestro, J.J., Miletić, S., Forstmann, B.U., 2019. Advances in techniques for imposing reciprocity in brain-behavior relations. Neurosci. Biobehav. Rev. 102, 327–336. https://doi.org/10.1016/j.neubiorev.2019.04.018

Turner, B.M., Sederberg, P.B., Brown, S.D., Steyvers, M., 2013. A method for efficiently sampling from distributions with correlated dimensions. Psychol. Methods 18, 368–384. https://doi.org/10.1037/a0032222

Turner, R., 2002. How much codex can a vein drain? Downstream dilution of activation-related cerebral blood oxygenation changes. Neuroimage 16, 1062–1067. https://doi.org/10.1006/nimg.2002.1082

Tustison, N.J., Avants, B.B., Cook, P.A., Yuanjie Zheng, Egan, A., Yushkevich, P.A., Gee, J.C., 2010. N4ITK: Improved N3 Bias Correction. IEEE Trans. Med. Imaging 29, 1310–1320. https://doi.org/10.1109/TMI.2010.2046908

Uǧurbil, K., Toth, L., Kim, D.S., 2003. How accurate is magnetic resonance imaging of brain function? Trends Neurosci. 26, 108–114. https://doi.org/10.1016/S0166-2236(02)00039-5

Ullsperger, M., Harsay, H.A., Wessel, J.R., Ridderinkhof, K.R., 2010. Conscious perception of errors and its relation to the anterior insula. Brain Struct. Funct. 214, 629–643. https://doi.org/10.1007/s00429-010-0261-1

Uludaǧ, K., Müller-Bierl, B., Uǧurbil, K., 2009. An integrative model for neuronal activity-induced signal changes for gradient and spin echo functional imaging. Neuroimage 48, 150–165. https://doi.org/10.1016/j.neuroimage.2009.05.051

Urai, A.E., Gee, J.W. De, Tsetsos, K., 2019. Choice history biases subsequent evidence accumulation. Elife 8, e46331. https://doi.org/10.7554/eLife.46331.001

van der Zwaag, W., Schäfer, A., Marques, J.P., Turner, R., Trampel, R., 2016. Recent applications of UHF-MRI in the study of human brain function and structure: a review. NMR Biomed. 29, 1274–1288. https://doi.org/10.1002/nbm.3275

van Maanen, L., Miletić, S., 2020. The interpretation of behavior-model correlations in unidentified cognitive models. Psychon. Bull. Rev. https://doi.org/10.3758/s13423-020-01783-y

van Wijk, B.C.M., Alkemade, A., Forstmann, B.U., 2020. Functional segregation and integration within the human subthalamic nucleus from a micro- and meso-level perspective. Cortex 131, 103–113. https://doi.org/10.1016/j.cortex.2020.07.004

Voss, A., Rothermund, K., Voss, J., 2004. Interpreting the parameters of the diffusion model: An empirical validation. Mem. Cognit. 32, 1206–1220. https://doi.org/10.3758/BF03196893

Wagenmakers, E.-J., Farrell, S., 2004. AIC model selection using Akaike weights. Psychon. Bull. Rev. 11, 192–196. https://doi.org/10.3758/BF03206482

Wang, S., Peterson, D.J., Gatenby, J.C., Li, W., Grabowski, T.J., Madhyastha, T.M., 2017. Evaluation of Field Map and Nonlinear Registration Methods for Correction of Susceptibility Artifacts in Diffusion MRI. Front. Neuroinform. 11. https://doi.org/10.3389/fninf.2017.00017

Weinberger, M., Mahant, N., Hutchison, W.D., Lozano, A.M., Moro, E., Hodaie, M., Lang, A.E., Dostrovsky, J.O., 2006. Beta Oscillatory Activity in the Subthalamic Nucleus and Its Relation to Dopaminergic Response in Parkinson's Disease. J. Neurophysiol. 96, 3248–3256. https://doi.org/10.1152/jn.00697.2006

Wessel, J.R., Jenkinson, N., Brittain, J.S., Voets, S.H.E.M., Aziz, T.Z., Aron, A.R., 2016. Surprise disrupts cognition via a fronto-basal ganglia suppressive mechanism. Nat. Commun. 7. https://doi.org/10.1038/ncomms11195

Wetzels, R., Wagenmakers, E.J., 2012. A default Bayesian hypothesis test for correlations and partial correlations. Psychon. Bull. Rev. 19, 1057–1064. https://doi.org/10.3758/s13423-012-0295-x

Woolrich, M.W., Behrens, T.E.J., Beckmann, C.F., Jenkinson, M., Smith, S.M., 2004. Multilevel linear modelling for FMRI group analysis using Bayesian inference. Neuroimage 21, 1732–1747. https://doi.org/10.1016/j.neuroimage.2003.12.023

Woolrich, M.W., Ripley, B.D., Brady, M., Smith, S.M., 2001. Temporal autocorrelation in univariate linear modeling of FMRI data. Neuroimage 14, 1370–1386. https://doi.org/10.1006/nimg.2001.0931

Worsley, K.J., 2001. Statistical analysis of activation images, in: Functional Magnetic Resonance Imaging. Oxford University Press, pp. 251–270. https://doi.org/10.1093/acprof:oso/9780192630711.003.0014

Zaidel, A., Spivak, A., Grieb, B., Bergman, H., Israel, Z., 2010. Subthalamic span of oscillations predicts deep brain stimulation efficacy for patients with Parkinson's disease. Brain 133, 2007–2021. https://doi.org/10.1093/brain/awq144

Zhang, Y., Brady, M., Smith, S., 2001. Segmentation of brain MR images through a hidden Markov random field model and the expectation-maximization algorithm. IEEE Trans. Med. Imaging 20, 45–57. https://doi.org/10.1109/42.906424

